# A network of RS splicing regulatory proteins controls light-dependent splicing and seedling development

**DOI:** 10.1101/2025.04.07.646831

**Authors:** Jennifer Saile, Hannah Walter, Moritz Denecke, Patrick Lederer, Laura Schütz, Andreas Hiltbrunner, Katharina Lepp, Sofia Lobato-Gil, Petra Beli, Andreas Wachter

**Affiliations:** Institute for Molecular Physiology (imP), University of Mainz, Hanns-Dieter-Hüsch-Weg 17, 55128 Mainz, Germany; Institute of Biology II, University of Freiburg, 79104 Freiburg, Germany and Signaling Research Centres BIOSS and CIBSS, 79104 Freiburg, Germany; Institute of Molecular Biology (IMB), Ackermannweg 4, 55128 Mainz, Germany; Institute of Developmental Biology and Neurobiology (IDN), Johannes Gutenberg-Universität, Hanns-Dieter-Hüsch-Weg 15, 55128 Mainz, Germany

**Keywords:** Alternative splicing, SR protein, light, photomorphogenesis, skotomorphogenesis

## Abstract

The light-induced change from skoto- to photomorphogenesis is a key switch in plant development that requires global transcriptome reprogramming. Earlier studies in *Arabidopsis thaliana* and other plant species have revealed the eminent role of alternative precursor mRNA splicing (AS), which allows fine-tuning the expression of numerous genes including light signaling and photosynthesis-related components in response to the ambient light conditions. Starting from the previous finding that AS changes induced by either light or metabolic signals are linked to phospho-signaling, we applied phospho-proteomics to identify proteins that undergo rapid changes in their phosphorylation status upon exposing etiolated seedlings to either light or sucrose. This approach revealed hyperphosphorylation of RS41, a member of the RS subfamily of serine/arginine-rich (SR) proteins. To study the function of the four RS genes *RS31*, *RS31a*, *RS40*, and *RS41*, a comprehensive set of single and higher order mutants was generated. A complete loss of RS function in the quadruple mutant caused sterility. Moreover, the important role of the RS proteins in seedling photomorphogenesis was demonstrated, with both redundant and specific functions in the regulation of hypocotyl elongation and cotyledon opening. We further identified the critical contribution of the RS proteins to light-dependent alternative splicing, being part of an intricate network of splicing regulatory components. Our study provides novel insight into the complex network of RNA-binding proteins that allow balancing light-responsive splicing and development in Arabidopsis seedlings.

## Introduction

Plants have a diverse set of light sensory and signaling systems that allow adjusting their biochemical activities, physiology, and development to this critical and fluctuating environmental factor. Accordingly, the early and critical phase of seedling establishment is highly responsive towards the ambient light conditions, balancing root and shoot growth to optimize nutrient uptake and light utilization for photosynthesis. Skoto- and photomorphogenesis refer to the developmental programs of seedlings grown in darkness and light, respectively, and differ amongst other parameters in root and hypocotyl growth, pigment accumulation, and cotyledon opening (Gommers and Monte, 2018). When etiolated seedlings are exposed to light, they undergo a rapid switch from skoto- to photomorphogenesis and photosynthesis is initiated. The light signals are perceived by a sophisticated set of photoreceptors (Galvão and Fankhauser, 2015; Podolec and Ulm, 2018), which elicit downstream signaling processes to reprogram gene expression and thereby adjust seedling development.

Light-dependent changes in gene expression occur at various levels, including alternative precursor mRNA splicing (AS) (Kathare and Huq, 2021) that allows generation of splicing variants from one gene by variable exon/intron definition. Transcriptome analyses of etiolated *Arabidopsis thaliana* seedlings demonstrated that light exposure alters the AS patterns of a large number of genes (Hartmann et al., 2016; Shikata et al., 2014). Comparing the AS output between wild type (WT) and phytochrome (phy) mutants, which have defects in red light sensing, indicated an involvement of phy signaling in light-dependent splicing, at least for some AS events and under certain light conditions in Arabidopsis seedlings (Shikata et al., 2014; Hartmann et al., 2016) and also in the protonema of *Physcomitrella patens* (Wu et al., 2014). Interestingly, exposing etiolated seedlings to external sugar provoked for several genes similar AS changes as observed upon illumination, suggesting a role for metabolic signaling in this process (Hartmann et al., 2016). Moreover, the observation of red light-induced AS in Arabidopsis seedlings mutated in all five *PHY* genes provided evidence that these splicing shifts can also occur independently of phy signaling (Mancini et al., 2016; Careno et al., 2023). Studying light-dependent AS of the Arabidopsis *RS31* gene, which encodes a member from the family of Serine/Arginine-rich (SR) proteins (Barta et al., 2010), suggested that a chloroplast-derived retrograde signal acts upstream of the regulation of this splicing event (Petrillo et al., 2014). Subsequent studies identified the central energy sensor kinases TARGET OF RAPAMYCIN (TOR) and SNF1-RELATED KINASE 1 (SnRK1) as key regulators of light-responsive AS in Arabidopsis seedlings (Riegler et al., 2021; Saile et al., 2023).

The observation of AS re-programming in response to changing light and metabolic conditions also sparked interest in the downstream regulators of this process. In general, the AS outcome is defined by the interplay of *cis*-regulatory elements within the pre-mRNA and *trans*-acting splicing regulators, including the aforementioned SR proteins and other classes of RNA-binding proteins (RBPs) (Staiger and Brown, 2013; Reddy et al., 2013). In a mutant screen for factors suppressing red light-dependent hypocotyl expansion in Arabidopsis, Shikata et al. (2012) identified the gene *REDUCED RED-LIGHT RESPONSES IN CRY1CRY2 BACKGROUND 1 (RRC1). RRC1* encodes a protein composed of an RNA binding domain and several regions that are predicted to be structurally disordered.

Hypocotyl elongation of *rrc1* mutants in red light was increased (Shikata et al., 2012), in line with the protein’s function as a positive regulator of photomorphogenesis that normally suppresses hypocotyl growth in response to light. This study also revealed altered AS patterns for several *SR* genes in an allelic series of *rrc1* mutants. Similar to *RRC1*, *SPLICING FACTOR FOR PHYTOCHROME SIGNALING* (*SFPS*) resulted from a mutagenesis screen for seedlings with elongated hypocotyls in red light (Xin et al., 2017). SFPS was shown to interact with phyB and co-localization of the two proteins in light-induced nuclear speckles was observed. Transcriptome-wide AS studies in *sfps* mutants revealed many splicing alterations compared to the WT, including light-regulated AS events. Interestingly, a following screen for SFPS-interacting proteins also identified RRC1 (Xin et al., 2019), underlining its critical role in light regulation. It was demonstrated that SFPS and RRC1 do not only co-localize with phyB in light-induced nuclear speckles, but also have common functions in light-regulated AS. The same screen uncovered another *bona fide* RBP interacting with SFPS, namely SWAP1 (SUPPRESSOR-OF-WHITE-APRICOTE/SURP RNA-BINDING DOMAIN-CONTAINING PROTEIN1), which can form a ternary complex with SFPS and RRC1 (Kathare et al., 2022). Single and double mutants in these factors showed a comparable extent of hyposensitivity to red light, suggesting that they can act together and/or on the same targets. Furthermore, a common set of regulated AS events was detected, pointing to an important function of the SWAP1-SFPS-RRC1 complex in light-dependent AS in Arabidopsis seedlings. In contrast to these positive regulators of photomorphogenesis, SWELLMAP2 (SMP2) was reported to antagonize phyB signaling in Arabidopsis with light-grown *smp2* mutants displaying shortened hypocotyls (Yan et al., 2022). In this study, an interaction between SMP2 and phyB was observed both in a yeast two hybrid assay and *in planta*. Moreover, a YFP-SMP2 fusion localized in the nucleoplasm and phyB-containing nuclear speckles in the hypocotyl from transgenic Arabidopsis seedlings grown under continuous red light. SMP2 is a homolog of the splicing factor SLU7, which was previously shown to be required for 3’ splice site selection (Frank and Guthrie, 1992), and Yan et al. (2022) described changes in the AS output of the circadian clock gene *RVE8* in the *smp2* mutant. Further studies are needed to examine whether SMP2 can also contribute to the regulation of light-dependent AS events and thereby may antagonize the functions of SWAP1, SFPS, and RRC1 in splicing control. Besides these studies in *A. thaliana*, deeper insights into the mechanisms of light-regulated AS were also gained in the moss *P. patens*. Screening for phy-interacting proteins identified HETEROGENEOUS NUCLEAR RIBONUCLEOPROTEIN H1 (HNRNP-H1) (Shih et al., 2019) and HNRNP-F1 (Lin et al., 2020), both of which regulate intron retention in a red light-dependent manner in *P. patens*.

The identification of different regulators of light-dependent AS in Arabidopsis and *P. patens* may reflect species-specific variation or indicate the involvement of an even larger number of RBPs. Accordingly, additional splicing regulatory proteins might contribute to this process.

Starting from the previous observation of fast and kinase-dependent AS changes, we searched in this study for RBPs that undergo rapid phosphorylation upon light and sucrose treatment in etiolated Arabidopsis seedlings and therefore may function in the downstream signaling. This approach resulted in the identification of RS41, an SR protein belonging to the plant-specific RS subfamily which is defined by the presence of a C-terminal RS domain enriched in RS dipeptides, and contains the additional members RS31, RS31a, and RS40. We observed that illumination can alter total transcript levels and/or AS of all four *RS* genes in etiolated seedlings, further suggesting their involvement in the light response. To investigate the functions of the *RS* genes, we generated via CRISPR/Cas9 a comprehensive set of single and higher order *rs* mutants. As a complete loss of *RS* functions in a quadruple mutant resulted in male sterility, we also established an inducible knockdown of *RS31* in the background of a double or triple knockout of the other *RS* genes. Examining these mutants’ responses to light revealed that the *RS* genes are regulators of AS and development in etiolated seedlings. Interestingly, *RS31* antagonized the effect of the other three *RS* genes in light-dependent suppression of hypocotyl elongation, indicating functional specialization within this group of splicing regulatory proteins. Furthermore, we demonstrated intricate and AS-based cross-regulation between the *RS* genes and *RRC1*, which has previously been shown to control light-dependent splicing and hypocotyl elongation. Our study reveals how a network of *RS* and other RBP genes functions in balancing light-dependent AS and seedling development in the transition from skoto- to photomorphogenesis.

## Results

### Phosphorylation of RS proteins can be altered upon light and sugar exposure of etiolated seedlings

Exposing etiolated Arabidopsis seedlings to either light or sugar can trigger similar AS changes in a set of genes (Hartmann et al., 2016). The rapidness of these AS responses and the observation that kinase inhibition can cause comparable AS shifts suggested the involvement of phospho-signaling in this process (Hartmann et al., 2016). This was further corroborated by the demonstration that the two central energy sensor kinases SnRK1 and TOR can affect light-regulated AS and seedling development (Riegler et al., 2021; Saile et al., 2023). To identify potential downstream targets of kinase signaling, we performed a phospho- proteome analysis using 6-d-old etiolated Arabidopsis wild type (WT) seedlings. The dark-grown seedlings were either illuminated with white light or exposed to exogenous sucrose for 30 min, while seedlings that were further kept in darkness served as control. Upon protein extraction from the seedling samples, phospho-peptides were enriched and analyzed via mass spectrometry. For the light-treated samples, the most pronounced change detected among the significantly altered phospho-peptides was a hyperphosphorylated serine residue within RS41 (Supplementary Fig. S1A; Supplementary Data Set. S1). The same serine residue in RS41 showed increased phosphorylation in response to sucrose treatment. From two independent phospho-proteome experiments, we were able to detect a total of six phospho-sites in RS41 and also four phosphorylated serine residues in its homolog RS40 (Supplementary Fig. S1B). In line with our findings for etiolated seedlings, sucrose treatment of Arabidopsis roots altered the phosphorylation status of several serine residues in RS40 and RS41 (Wu et al., 2019). Inspection of their overall phosphorylation potential based on the entries in the plant PTM viewer database (Willems et al., 2019, Supplementary Fig. S1B) confirmed the phospho-sites detected by us and revealed numerous additional sites, all being located within the RS domains of RS40 and RS41. The RS subfamily from Arabidopsis comprises two additional members, RS31 and RS31a. Multiple phospho-sites within their RS domains and, in case of RS31, also two phosphorylated serine residues within one of the two RNA recognition motifs (RRMs) are deposited in the plant PTM viewer database. Accordingly, the RS proteins can be phosphorylated at multiple sites and under various conditions. Moreover, the altered phosphorylation of RS41 upon light and sugar exposure in etiolated seedlings suggested its possible involvement in regulating the changes in AS and seedling development under those conditions.

Previous studies identified TOR and SnRK1 as upstream regulators of light-mediated AS events (Saile et al., 2023) and revealed reduced phosphorylation of RS40 and RS41 upon TOR inactivation in light-grown seedlings (Scarpin et al., 2020). Therefore, we hypothesized that the kinases might control AS by modulating RS protein activity due to their altered phosphorylation in response to metabolic cues. To study the potential impact of TOR on RS41 protein levels in etiolated seedlings, we generated a transgenic line containing a tagged RS41 version (*35S:RS41-HA_3_*) for immunodetection, along with an estradiol-inducible expression cassette of an artificial microRNA (*i-amiR*) for knocking down *TOR*, as previously reported (Saile et al., 2023). Exposing these seedlings to estradiol strongly reduced *TOR* transcript level in two independent lines, confirming efficient *TOR* knockdown (Supplementary Fig. S2A). Analyzing RS41-HA_3_ protein levels via immunoblot detection revealed for the estradiol-treated seedlings an ∼1.5-fold increase in RS41-HA_3_ relative to the mock sample (Supplementary Fig. S2B, compare 0 min samples). To test whether steady state levels of RS41-HA_3_ are altered in response to metabolic and light signals and if this might depend on TOR, we treated dark-grown seedlings of this double transgenic line with sucrose or white light in either the absence or presence of estradiol. Neither sucrose nor light exposure resulted in a significant change in RS41-HA_3_ levels (Supplementary Fig. S2B), however, in line with the finding for the 0 min samples, all of the *TOR* knockdown samples showed an elevated level of the RS41 fusion protein compared to the corresponding mock samples. Given our previous observation that knocking down *SnRK1* similarly affects light-dependent AS and seedling development as seen upon *TOR* knockdown (Saile et al., 2023), we also generated a double transgenic line carrying *35S:RS41-HA_3_* in an *i-amiR-SnRK1* background. Again, we observed slightly but consistently increased RS41-HA_3_ levels upon kinase knockdown, independent of the light condition (Supplementary Fig. S2C). Moreover, the absence of light-induced changes in RS41-HA_3_ protein levels was also confirmed in a transgenic line without the *i-amiR* construct (Supplementary Fig. S2D). In summary, we observed that RS41 protein levels can be negatively regulated by both SnRK1 and TOR in etiolated seedlings, and further studies are required to elucidate the underlying mechanism.

### Expression of *RS* genes is regulated by light

To examine whether the expression of the *RS* genes is controlled by light, we first analyzed their total transcript levels in 6-d-old etiolated WT seedlings that were exposed to either white, red, far-red or blue light for 6 h compared to non-illuminated samples. RT-qPCR revealed diminished transcript levels of *RS31* and *RS40* in the light samples, whereas *RS31a* and *RS41* levels remained overall unaffected by the light treatments (Fig. 1A-D). These findings were in agreement with the RNA-seq data from a previous study (Hartmann et al., 2016) (Supplementary Fig. S3A-D). Moreover, comparison of the transcript levels based on the TPM values from those RNA-seq data demonstrated that steady state transcript amounts of *RS31*, *RS40,* and *RS41* were in a similar range, whereas *RS31a* transcripts were more than ten-fold less abundant in this stage of Arabidopsis seedling development.

**Fig. 1.**
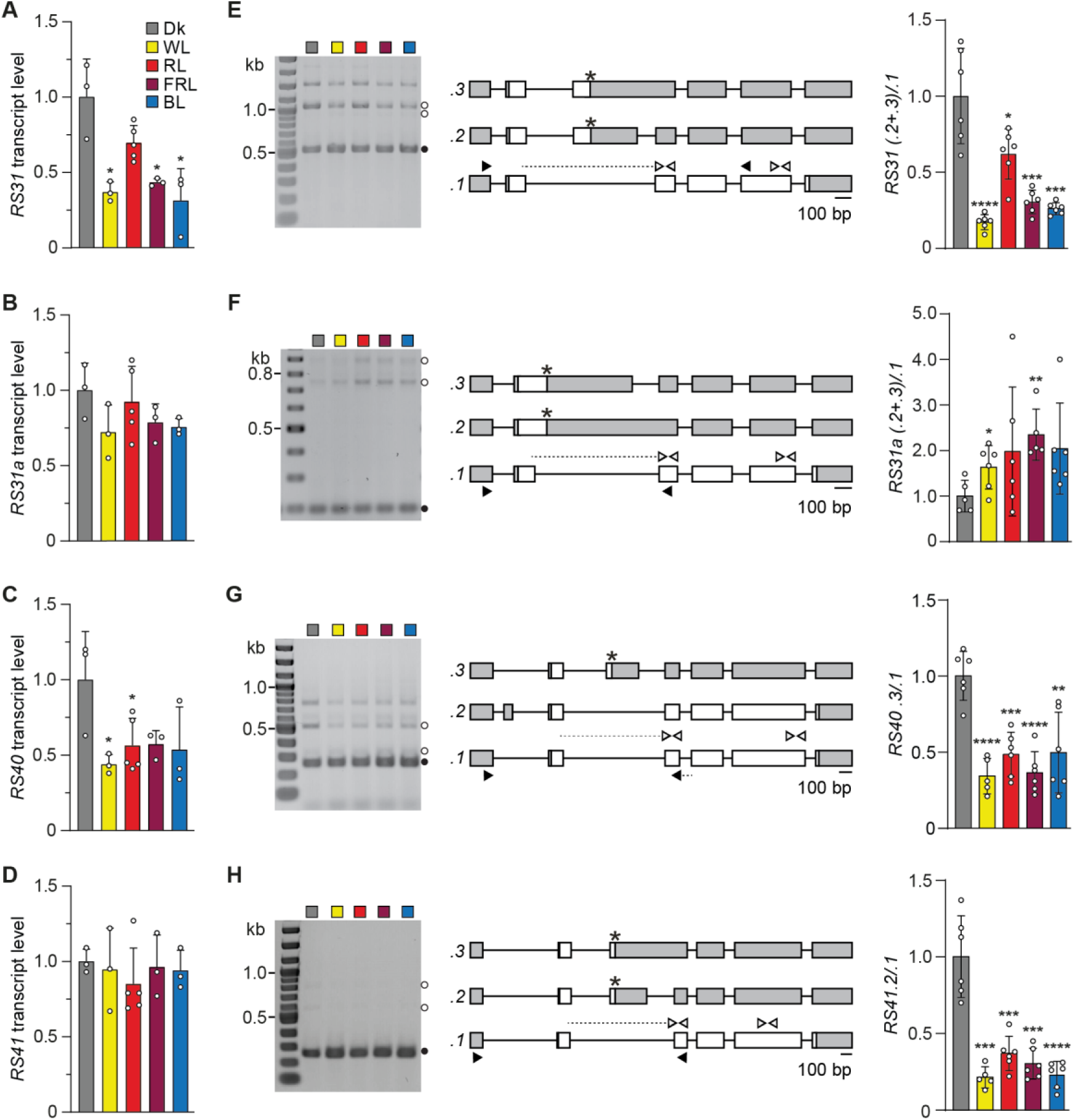
Light alters transcript levels and AS of several *RS* genes. **A – D)** Total transcript levels of *RS31*, *RS31a*, *RS40,* and *RS41* relative to the reference transcript *PP2A* in 6-d-old etiolated WT seedlings that were either kept in darkness (Dk) or exposed to white light (WL, ∼10 μmol m^-2^ s^-1^), red light (RL, ∼8 μmol m^-2^ s^-1^), far-red light (FRL; ∼6 μmol m^-2^ s^-1^), or blue light (BL∼8 μmol m^-2^ s^-1^) for 6 h. Bars represent mean values (n = 3 to 5; individual data points are shown as dots) +SD, normalized to mean value in dark. Statistical significance was determined by two-tailed Student’s t-test against corresponding dark control (P value: *P < 0.05). **E – H)** AS patterns of *RS31*, *RS31a*, *RS40,* and *RS41* from samples as described above that were either kept in darkness or illuminated (WL ∼10 μmol m^-2^ s^-1^, RL ∼8 – 11 μmol m^-2^ s^-1^, FRL ∼6 – 11 μmol m^-2^ s^-1^, BL ∼8 – 11 μmol m^-2^ s^-1^). (Left) Representative agarose gel pictures of RT-PCR products. Filled and open circles indicate representative and alternative variants, respectively, confirmed by sequencing. DNA size marker in first lane each consisted of DNA fragments in 100 bp increments from 100 bp to 1 kb, followed by 1.2 kb, 1.5 kb, and 2 kb. Top most signal for *RS31* corresponds to length of variant containing all introns within the amplified sequence. Top most signal for *RS40* could not be identified and was undetectable using a Bioanalyzer. (Middle) Transcript models with introns, coding exons, and UTRs shown as lines, white boxes, and grey boxes, respectively. Premature termination codons indicated by asterisks. Primer pairs used for RT-qPCR and co-amplification PCR, respectively, are depicted by white and black arrowheads. Primer with dotted line indicates primer binding site at a splicing junction. Numbering of variants is for the major coding variant according to .*1* from TAIR 10; the other variants are consecutively numbered in the order of their size. Start of first and end of last exon have been set according to representative variant *.1*. (Right) Quantification of the splice isoform ratios. Data represents mean ± SD, normalized to the mean dark control, with individual data points shown as dots (n = 5 to 6). A two-tailed student’s t-test was performed to analyze significant differences compared to the dark control (P values: *P < 0.05, **P < 0.01, ***P < 0.001, ****P < 0.0001).

The expression of many RBPs is regulated on the level of AS, typically resulting in the increased formation of unproductive variants at the expense of the coding variant as part of negative auto- and cross-regulatory feedback loops. Accordingly, light-regulated AS has previously been reported for several *RS* genes (e.g. Shikata et al., 2014; Petrillo et al., 2014). Therefore, we inspected the AS patterns of the four *RS* genes in the etiolated seedlings kept in darkness or illuminated with the different light qualities. We detected for all four *RS* genes two or more splicing variants, which were identified by sequencing (Fig. 1E-H, Supplementary Fig. S3E-G). Interestingly, pronounced light-induced AS shifts were seen for *RS31*, *RS40*, and *RS41*, all of which showed a relative decrease of the unproductive variants upon illumination. Given that the *RS31* and *RS40* genes also showed reduced levels of total transcripts in response to light treatment, the corresponding AS changes may be the consequence of a diminished auto-regulation. In contrast, total *RS41* transcript amounts were similar in the dark and light conditions. Therefore, this altered splicing pattern may be the consequence of a reduced level of negative cross-regulation by *RS31* and *RS40* or other RBPs. Finally, an opposite trend in the AS response was seen for *RS31a,* with a slight increase in the relative levels of unproductive isoforms upon illumination. Taken together, the expression of the *RS* genes is highly responsive to light treatment in etiolated seedlings, and the response patterns indicate the existence of interconnected and gene-specific regulatory mechanisms.

### An *rs* quadruple mutant shows reduced fertility

Our experiments revealed regulation of the *RS* genes during the light response of etiolated seedlings and we speculated that the individual *RS* genes may have specific and common functions in this and possibly other processes. To further study the role of the RS proteins, we used the CRISPR/Cas9 system to generate *rs* single and higher order knockout mutants. Sequencing analysis of the mutants identified single nucleotide insertions or deletions within the *RS* genes, resulting in frameshifts that cause premature termination codons (PTCs) downstream and in vicinity of the mutation sites (Fig. 2). Due to the early positioning of the corresponding PTCs relative to the regular stop codon, these mRNAs are likely unproductive and the plants were considered as *bona fide* knockout mutants in the respective *RS* genes. Triple and quadruple *rs* mutants were generated by transforming the *rs40 rs41_C_* double mutant with the CRISPR/Cas9 construct targeting *RS31* and *RS31a* and are hereafter referred to as *rsT_C_* (*rs31a rs40 rs41_C_)* and *rsQ_C_* (*rs31 rs31a rs40 rs41_C_*), respectively (Fig. 2). When growing *rsQ_C_* plants to the stage of seed setting we noted that the loss of all four *RS* genes resulted in stunted siliques and a strongly reduced seed yield (Supplementary Fig. S4A-D). This observation is in contrast to a previous study (Yan et al., 2017) reporting no morphological abnormality in a quadruple mutant that carries T-DNA insertions in all four *RS* genes. Residual RS activity deriving from one or several of the T-DNA-containing mutant alleles in the respective mutant may account for this difference. To determine whether the loss of individual *RS* genes can already affect the seed yield, we also measured the silique sizes in WT, *rs* double, and *rsT_C_* mutants. Interestingly, the concomitant loss of *RS40* and *RS41* also reduced the silique length, while the *rs31 rs31a_C_* double mutant did not significantly differ in this trait from the WT (Supplementary Fig. S4C, D). Moreover, knocking out *RS31a* besides *RS40* and *RS41* in the *rsT_C_* had no additional effect compared to the *rs40 rs41_C_* double mutant, whereas much shorter siliques were found in *rsQ_C_* (Supplementary Fig. S4C, D). Siliques of the *rsQ_C_* mutants were not only much shorter, but also contained mainly aborted and shrunken seeds (Supplementary Fig. S4C). Moreover, the anthers of the quadruple mutant exhibited a strongly reduced amount of pollen compared to WT plants and were mis-formed (Supplementary Fig. S4E). To test whether the reduced seed yield of the *rsQ_C_* mutant is caused by a pollen defect, we pollinated stigmas of *rsQ_C_* mutants with WT pollen and also performed the reciprocal crossing experiment. We observed that mutant pollen failed to result in seed setting in WT mother plants, whereas the transfer of WT pollen to *rsQ_C_* stigmas generated WT-like siliques (Supplementary Fig. S4F, G). These data suggest that the impaired seed production of the *rsQ_C_* mutant is due to male sterility. Moreover, comparison of the silique phenotypes in two types of double, one triple, and the quadruple mutants provided evidence that the RS proteins have specific and redundant functions in this context.

**Fig. 2.**
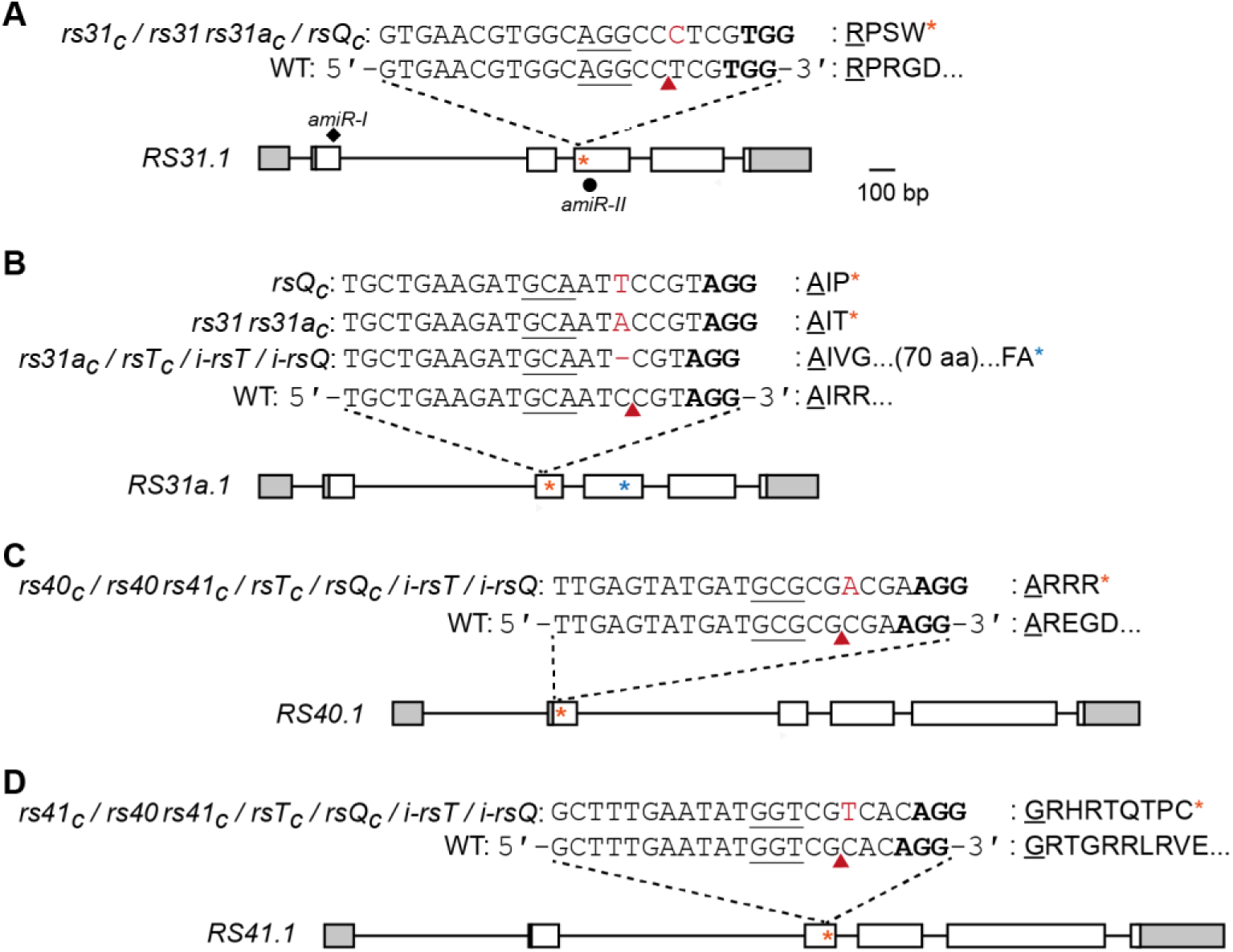
Models of *RS* genes with CRISPR-induced mutations. Representative transcript models of *RS31* (**A**), *RS31a* (**B**), *RS40* (**C**), and *RS41* (**D**), with introns, coding exons, and UTRs shown as lines, white boxes, and grey boxes, respectively. The target sites of the two inducibly expressed *amiRs (I/II)* against *RS31* are shown by a black rhomb and circle, respectively. The positions of the sgRNA target sites are indicated by dashed lines and sgRNA target sequences are shown above the transcript models, with the bold sequence showing the PAM. Cas9 cleavage sites are indicated by the red arrowhead and corresponding mutations (insertion or deletion) are marked in red, followed by the corresponding amino acid (aa) sequence on the right. The displayed aa sequence starts with the codon triplet prior the cleavage site (underlined). Premature termination codons are generated by the mutation-induced frameshift and depicted as asterisks (orange/blue). Drawn to scale.

To overcome the problem of male sterility in *rsQ_C_*, we generated *i-amiR* constructs to specifically downregulate *RS31* in an estradiol-inducible manner in the CRISPR-based knockout background of the other *RS* genes (Fig. 2). Two independent *i-amiR-RS31* constructs for targeting different sequences of the *RS31* transcript were generated and used for transformation of either *rs40 rs41_C_* double or *rsT_C_* mutants, resulting in *i-rsT-I/II* and *i-rsQ-I/II*, respectively. Corresponding plants showed normal plant development including regular silique generation when grown without estradiol, proving sufficiently tight control of *amiR* production (Supplementary Fig. S5A). To determine the efficacy of our knockdown approach, we exposed etiolated *i-rsT* and *i-rsQ* seedlings to estradiol for 3 d and analyzed *RS31.1* expression in several independent lines for each construct and genetic background. RT-qPCR analysis demonstrated that estradiol treatment strongly reduced *RS31.1* transcript levels in both *i-rsT (-I/II)* lines (Supplementary Fig. S5B). When comparing the mock-treated samples, *RS31.1* levels were in general higher in the mutant lines than in the WT. As the non-induced *i-rsT* corresponds to an *rs40 rs41* double mutant, a negative cross-regulation of *RS31* by RS40 and/or RS41 can be concluded from this data. Similar observations were made for the *i-rsQ-I/II* lines (Supplementary Fig. S5C). Accordingly, the *amiRs* were functional and their induction resulted in a triple or quadruple *rs* mutant, which allowed us to further study the role of the *RS* genes in seedling development.

### Hypocotyl elongation under red light is reduced upon *RS31* knockdown, while loss of the other *RS* genes has the opposite effect

To examine a possible role of the *RS* genes in skoto- and photomorphogenesis, we analyzed hypocotyl lengths of the *rs* mutants grown in darkness or under different light qualities. Estradiol-induced knockdown of *RS31* caused a pronounced shortening of the hypocotyls in darkness for both the *i-rsT* and *i-rsQ* lines (Fig. 3A, B; Supplementary Fig. S6A, B), while WT seedlings showed as expected no response to the estradiol treatment. Intriguingly, an even more pronounced reduction in hypocotyl lengths upon estradiol treatment was seen for the *i-rsT* and *i-rsQ* mutants when grown under red light conditions. Thus, the downregulation of *RS31* in these mutants resulted in red light hypersensitivity, uncovering a role of *RS31* as negative regulator of the corresponding light response. The lines carrying *amiR-II* showed a stronger response than those containing the *amiR-I*, being in line with a slightly more pronounced reduction in *RS31* levels (Fig. 5A). Moreover, miRs can also inhibit translation; therefore, differences in their efficiency degree might be even higher than estimated based on the levels of the target transcript. When comparing the dark-normalized mean hypocotyl lengths of the mock-treated WT, *i-rsT*, and *i-rsQ* seedlings under continuous red light (Fig. 3C, Supplementary Fig. S6C), we noted that both mutants had longer hypocotyls and therefore showed a diminished light response. This effect was most pronounced in the mock-treated *i-rsQ* mutant, which lacks besides *RS40* and *RS41* also the *RS31a* gene. Thus, all of these three *RS* genes normally function as suppressors of hypocotyl elongation and thereby activate photomorphogenesis. The estradiol-treated *i-rsT* mutant showed WT-like hypocotyl lengths (Fig. 3C), indicating that the loss of two suppressors (*RS40*, *RS41*) and one activator (*RS31*) of hypocotyl elongation has a balancing effect. In contrast, the induced *i-rsQ* mutant had slightly longer hypocotyls than the WT, suggesting that the simultaneous loss of *RS40*, *RS41*, *RS31a* cannot be fully compensated by the knockdown of *RS31*.

**Fig. 3.**
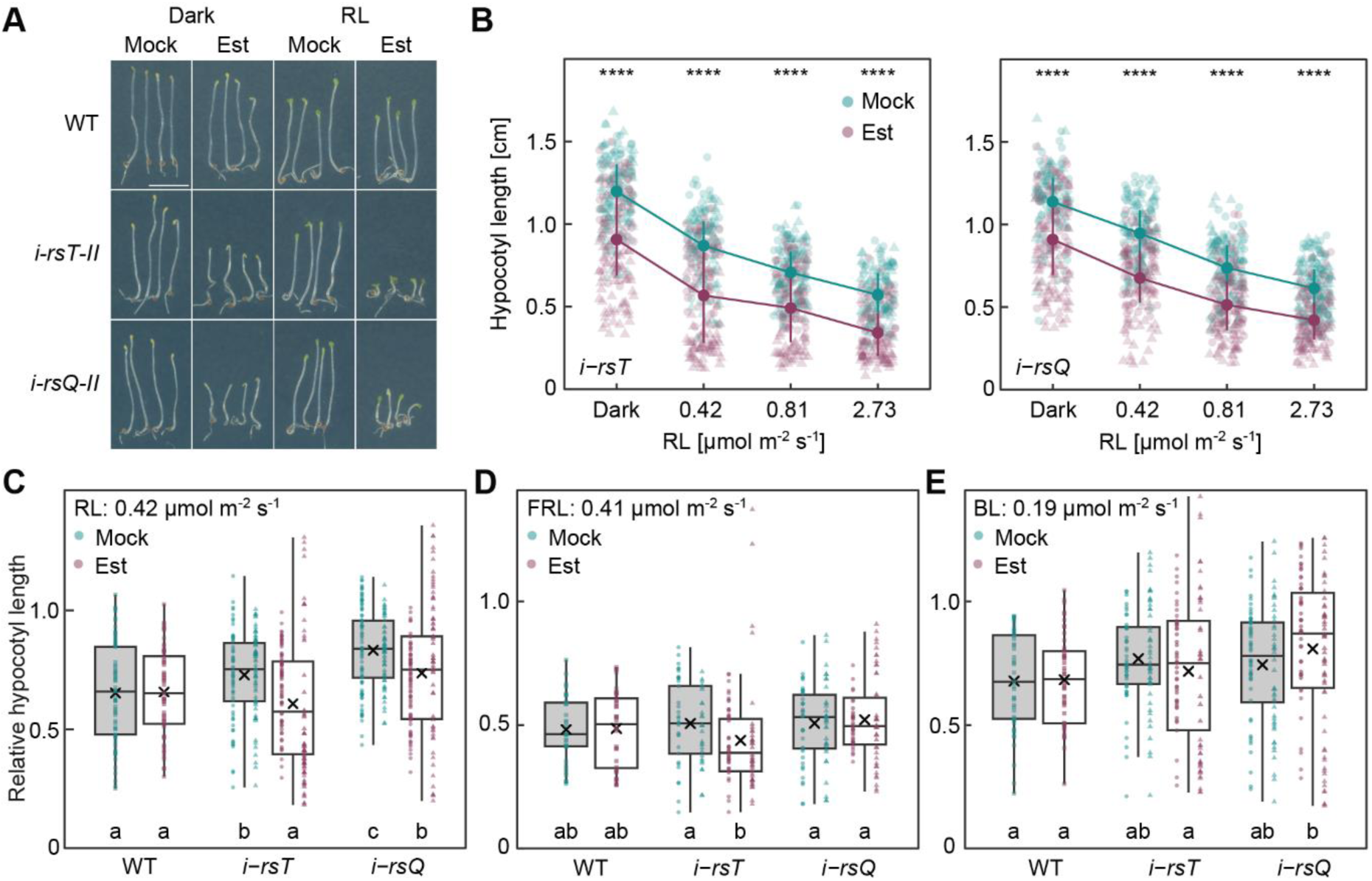
*RS31* acts a positive regulator of hypocotyl elongation under red light, whereas the other *RS* genes have the opposite function. **A)** Representative pictures of WT, *i-rsT-II* (*i-amiR-RS31-II* in *rs40 rs41_C_*), and *i-rsQ-II* (*i-amiR-RS31-II* in *rs31a rs40 rs41_C_*) seedlings grown for 4 d in darkness or red light (RL, 0.42 µmol m^-2^ s^-1^) under mock or Estradiol (Est) treatment. White scale bar corresponds to 0.5 cm. **B - E)** Absolute (B) and relative (C - E) hypocotyl length of WT, *i-rsT*, and *i-rsQ* grown under indicated light qualities and intensities or in darkness on plates containing either mock or Est. Circles and triangles indicate two lines with independent *amiR-RS31* constructs (I and II) which were used. For the absolute hypocotyl lengths (B), interquartile range and mean are depicted as vertical line and circle, respectively, and asterisks indicate significant differences of the comparison between Est and mock samples based on Kruskal-Wallis test with Dunn’s *post-hoc* test (α = 0.05) and ****P ≤ 0.0001 (n = 117 to 153; individual data points shown as dots). In (C - E), values were normalized to average length of dark-grown seedlings for each genotype, and interquartile range, maximum and minimum, median, and mean values are depicted as box, whiskers, middle line, and cross, respectively (n = 33 to 82; individual data points shown as dots). Letters a to c indicate significant group differences tested by Kruskal-Wallis test with Dunn’s *post-hoc* test (α = 0.05).

We next tested whether the *RS* genes also contribute to the development of seedlings grown under continuous far-red and blue light. In far-red light, the mock-treated *i-rsT* and *i-rsQ* lines had similar relative hypocotyl lengths as the WT (Fig. 3D), revealing that the loss of *RS40* and *RS41* in the double mutant and in addition *RS31a* in the triple mutant does not alter far-red light-dependent hypocotyl expansion. Knocking down *RS31* via amiR induction did not alter the hypocotyl length in the *i-rsQ* mutant. Slightly reduced hypocotyl lengths were seen upon Est treatment of *i-rsT* grown at 0.41 µmol m^-2^ s^-1^ far-red light, while no changes were detected at the other intensities analyzed (Supplementary Fig. S6D). Similarly, *rs* mutant seedlings grown in blue light showed no or only a minor increase in hypocotyl length in comparison of the WT and the non-induced mutants (Fig. 3E). Upon *RS31* knockdown, hypocotyl lengths were unchanged or, in case of *i-rsQ*, even slightly increased (Fig. 3E, Supplementary Fig. S6E). Taken together, the *RS* genes play an important and balancing role in red light-dependent hypocotyl elongation, but have little to no involvement in responses to far-red and blue light under the tested conditions.

Given that red light is sensed via phyB and previous studies demonstrated co-localization of splicing regulatory proteins with this photoreceptor in nuclear photobodies (Xin et al., 2019; Kathare et al., 2022), we addressed the question whether RS proteins can also localize in this subnuclear domain. Sub-cellular localization studies of RS41-GFP upon transient expression showed its nuclear localization, ranging from a more uniform distribution within the nucleoplasm to strong nuclear speckles in different cells and independent of whether the leaves were kept in darkness or light (Supplementary Fig. S7). In contrast, a co-transformed phyB-RFP construct localized in nuclear photobodies in case of the illuminated samples, being clearly distinct from the RS41-GFP localization pattern. Accordingly, at least a major proportion of RS41-GFP is not co-localizing with phyB under the tested conditions, and therefore may act independently of phyB and the aforementioned splicing regulators.

### RS overexpression can induce cotyledon opening in darkness

To gain further insights into *RS* gene functions, we also generated *RS* overexpression (OE) lines, in which the individual *RS* coding sequences were placed under control of the CaMV 35S promoter. Using RT-qPCR, we determined total *RS* transcript levels and selected the homozygous lines with highest steady state levels of the corresponding *RS* transcripts for our further analyses (Supplementary Fig. S8A). Compared to WT, the *RS31a* and *RS41 OE* lines showed an about 20-fold higher expression of the corresponding *RS* gene, whereas *RS31 OE* displayed a four-fold overexpression level (Supplementary Fig. S8A). For the *RS40 OE* line, an only ∼2-fold increase in *RS40* transcripts compared to WT was observed, and the same increase was seen in the *35S::EGFP* control line. Accordingly, this increase may be a consequence of the antibiotic selection performed for the *RS OE* and *EGFP* lines and therefore not result from expression of the transgene. Based on the low expression level of *RS31a* compared to the other *RS* genes in WT seedlings (Supplementary Fig. S3), we also performed RT-qPCR with a primer pair directed against a region in the *OCS* terminator that is shared between all constructs and thereby allows to compare the *RS OE* lines amongst each other. Analysing 10-d-old light grown and 6-d-old etiolated seedlings, the selected *RS41 OE* line was found to give the highest expression (Supplementary Fig. S8B, C). The other three constructs showed overall comparable levels of transcript accumulation, with the exception of higher *RS31* accumulation in the corresponding *OE* line for the etiolated but not the light-grown seedlings. These samples also differed in the age of the seedlings and a mannitol treatment of the etiolated seedlings. Accordingly, the variation in *RS31* levels may be condition-specific and result from post-transcriptional regulation. Taken together, the *RS OE* lines had overall similar levels of the respective *RS* genes and could be directly compared, except the *RS41 OE* line with a significantly higher expression level.

The *RS OE* lines were then grown under long day conditions and their development was scored (Supplementary Fig. S8D). A particularly striking phenotype was observed for the *RS41 OE* lines that showed dwarfism and downward curled rosette leaves (Supplementary Fig. S8E). In addition, *RS31 OE*, *RS40 OE,* and *RS41* OE plants showed an overall delayed development compared to WT, the *EGFP* vector control line, and the *RS31a OE* plants (Supplementary Fig. S8D, F).

When growing the *OE* lines in darkness, they exhibited increased cotyledon opening, in contrast to the mainly closed cotyledons observed in WT and *rsT_C_* seedlings (Fig. 4A-C). Measurement of the opening angle confirmed that *RS OE* seedlings had more widely opened cotyledons, with significant differences observed in the *RS31*, *RS31a*, and *RS41 OE* lines compared to WT and *rsT_C_* seedlings. It is worth noting that *RS41* overexpression caused the strongest phenotype, possibly due to the highest level of overexpression among the studied lines. Moreover, the weakest effect was seen for the *RS31* OE line, despite the second highest expression level among the *RS* OE lines in etiolated seedlings. This might again point to a distinct function of *RS31* as also seen in the context of hypocotyl elongation. In line with enhanced photomorphogenesis upon *RS41* overexpression, relative hypocotyl lengths of this mutant were strongly reduced compared to WT seedlings upon growth under continuous red and far-red light (Fig. 4D, E, Supplementary Fig. S9). A less pronounced but for some intensities still significant reduction of hypocotyl length in *RS41 OE* compared to WT was seen for seedlings grown under blue light (Fig. 4F). No significant change in hypocotyl length was observed for the other three *RS* OE lines when grown in the different light qualities (Fig. 4D-F, Supplementary Fig. S9). This observation may again be explained by specificity among the RS genes, but also be related to the different levels of overexpression.

**Fig. 4.**
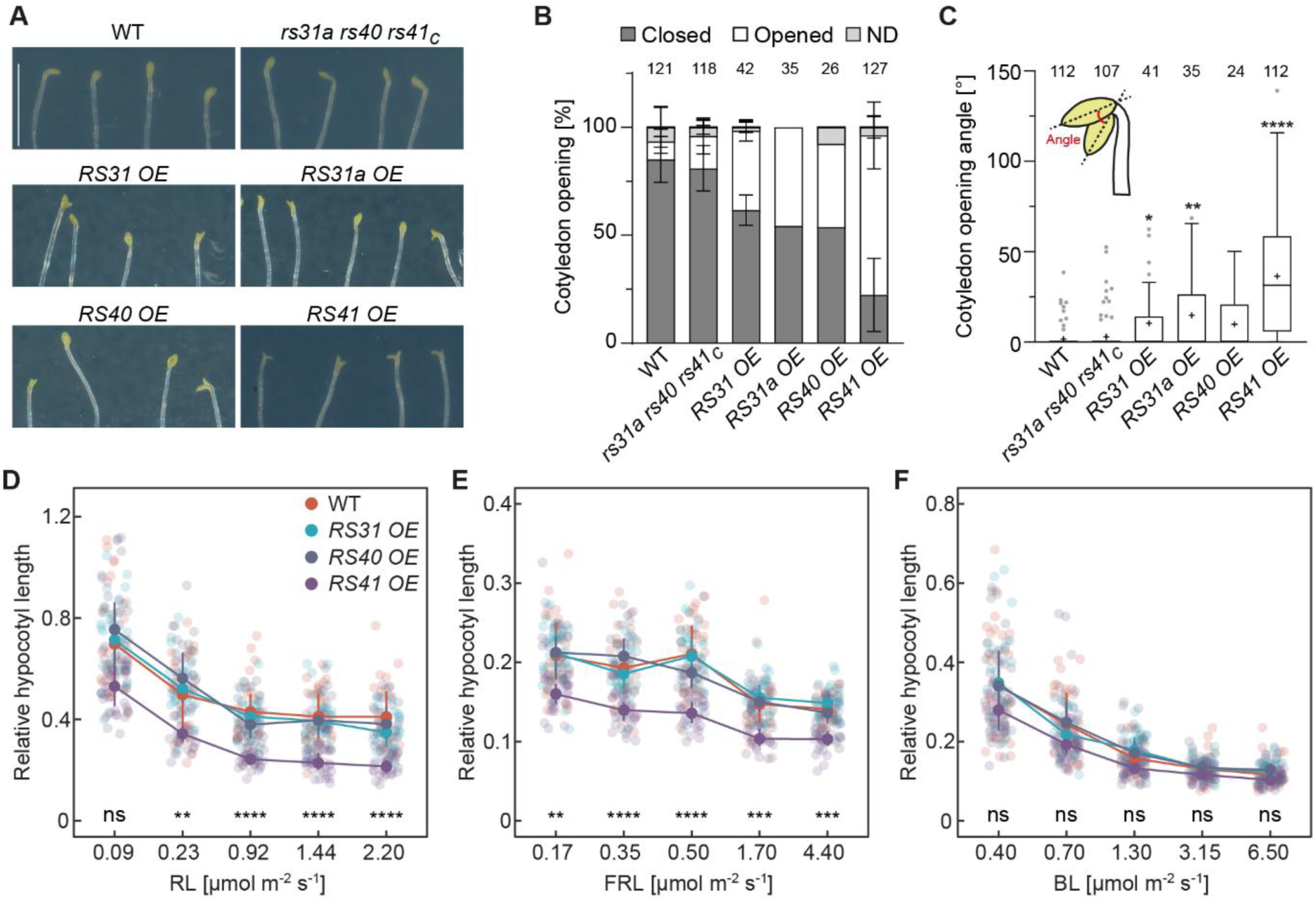
*RS41* overexpression causes cotyledon opening in darkness and hypocotyl shortening in light. **A)** Photographs of representative 6-d-old etiolated seedlings that were grown on ½ MS plates. White scale bar (applies to all images): 5 mm. **B)** Percentage of closed and opened cotyledons for 6-d-old etiolated seedlings. Displayed are mean values ± SD. Number of seedlings shown on top; these were derived from 1 to 4 independent experiments (*RS31a OE* and *RS40* OE: 1; *RS31* OE: 2; WT, *rs31a rs40 rs41_C_* and *RS41* OE: 4). **C)** Cotyledon opening angle in dark-grown seedlings. Growth conditions are as described in (A). Asterisks indicate significant differences to WT based on Kruskal-Wallis test with Dunn’s post-hoc test (P values: *P < 0.05, **P < 0.01, ***P < 0.001, ****P < 0.0001). Inset shows schematic visualization of cotyledon opening angle measurement in an etiolated seedling. **D – F)** Relative hypocotyl length of WT and *RS* OE lines grown for 4 d on plates under indicated continuous (D) red light (RL), (E) far-red light (FRL), or (F) blue light (BL) conditions. Values were normalized to average length of dark-grown seedlings for each genotype. Interquartile range and mean are depicted as vertical line and circle, respectively. Asterisks indicate significant differences of the comparison between WT and *RS41 OE* based on Kruskal-Wallis test with Dunn’s *post-hoc* test (α = 0.05) and ns > 0.05, *P ≤ 0.05, **P ≤ 0.01, ***P ≤ 0.001, ****P ≤ 0.0001 (n = 25 to 61; individual data points shown as dots).

### *RS* genes regulate light-dependent splicing events

Our experiments uncovered an important role of the *RS* genes in light-dependent seedling development and we hypothesized that this might involve the RS proteins’ function in AS regulation. To test this hypothesis, we grew the *i-rsT* and *i-rsQ* mutants in darkness followed by estradiol treatment to knock down *RS31* expression and examined the splicing patterns of several genes with light-regulated AS events. As already noted in course of the *i-amiR* line screening (Supplementary Fig. S5B, C), mock-treated samples of the *i-rsT* and *i-rsQ* mutants showed a ∼2-fold increase in *RS31* transcript levels in comparison to WT, indicating that the loss of *RS40* and/or *RS41* releases negative cross-regulation of *RS31* expression (Fig. 5A). Negative feedback control of *RS31* by *RS31a* seems not to occur under these conditions as *RS31* levels in mock-treated *i-rsQ* were not further increased compared to mock-treated *i-rsT*. Estradiol treatment for *amiR* induction in the mutants caused a strong reduction of *RS31* transcript levels, which were then back in the range of the WT or clearly below it (Fig. 5A). Accordingly, *amiR*-induced degradation of *RS31* not only compensates the upregulation seen in the *rs40 rs41_C_* background, but also leads to a further decrease of the target transcript level.

**Fig. 5.**
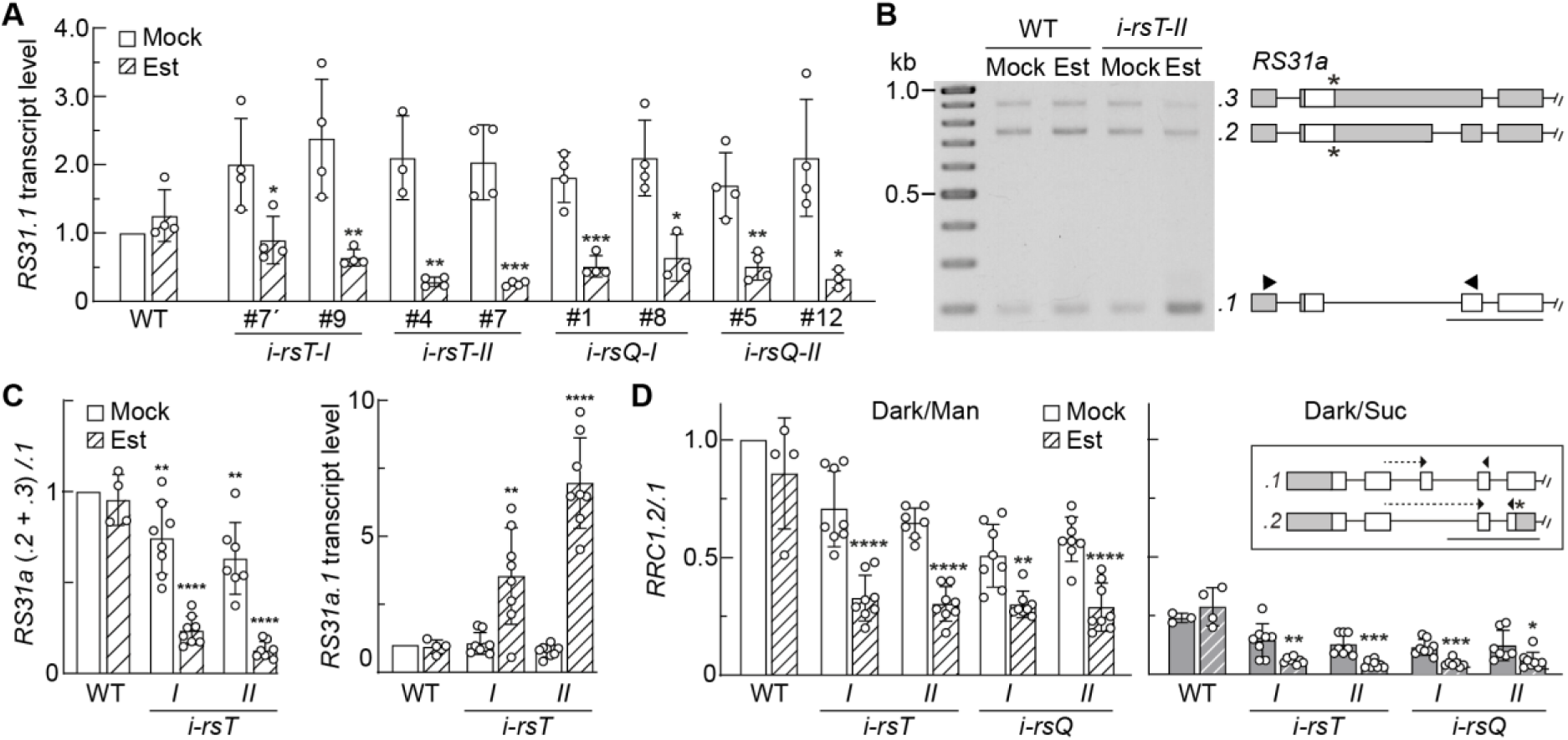
*RS* genes regulate light-dependent splicing events. **A)** Relative transcript level of *RS31.1* in 6-d-old etiolated seedlings, treated with either mock (white bars) or estradiol (Est, hatched bars) for 3 d. Displayed are mean values (n = 3 - 4; individual data points shown as dots) ± SD, normalized to the WT mock samples. Statistical significance was determined by two-tailed Student’s t-test against corresponding mock control (P values: *P < 0.05, **P < 0.01, ***P < 0.001) **B)** Agarose gel picture displaying *RS31a* splice isoforms upon co-amplification PCR. Plant material and growth conditions are as described in (A). DNA size ladder with 100 bp increments. Models of *RS31a* splicing variants are displayed next to the corresponding bands, with lines representing introns, grey and white boxes displaying UTRs and coding exons, respectively. Asterisk marks the position of a premature termination codon, and arrowheads depict binding sites of primers used for co-amplification PCR. Scale bar: 500 nts. **C)** (Left) Quantification of *RS31a* splice variant ratios via Bioanalyzer. Displayed are mean values (n = 4 - 8; individual data points as dots) ± SD, normalized to the WT mock sample. For the analysis, two sublines per *i-rsT* were combined. A one-sample t-test was performed in comparison to Col-0 Mock (P values: **P < 0.01, ****P < 0.0001). (Right) Relative transcript level of *RS31a.1* normalized to WT mock. Sample information and display features as described for part on the left. Statistical significance was determined by two-tailed Student’s t-test against corresponding mock control (P values: **P < 0.01, ***P < 0.001). **D)** *RRC1* AS ratio of 6-d-old etiolated seedlings, quantified via RT-qPCR. The seedlings were treated with either mock or estradiol (Est, hatched bars) for 3 d (left). Additionally, 6-d-old seedlings were treated with either 1.06% mannitol or 2% sucrose and then kept in darkness for 6 h (right). Displayed are mean values ± SD, normalized to the WT Mock samples (n = 3 – 8, with individual data points shown as dots; two independent *i-rsT* and *i-rsQ* sublines were combined each). A two-tailed Student’s t-test was performed in comparison to corresponding Mock control (P values: *P < 0.05, **P < 0.01, ***P < 0.001, ****P < 0.0001). AS event of *RRC1* is shown in the inset. Arrowheads mark binding sites of primers used for RT-qPCR.

As negative feedback regulation of RBPs often occurs on the level of AS, we next tested whether the splicing pattern of *RS31a* is altered upon knocking out/down the other members of the RS subfamily. In samples from the WT and the mock-treated *i-rsT* mutants, the PTC-containing variants *.2* and *.3* accumulated to a significant level besides the productive variant *RS31a.1* (Fig. 5B). Quantifying the ratio of unproductive to productive *RS31a* variants revealed a slight reduction in the mock-treated *i-rsT* lines compared to WT (Fig. 5C, left), suggesting that the loss of *RS40* and/or *RS41* results in a partial release of negative feedback control of *RS31a* expression, similar as observed for *RS31*. Estradiol-induced knockdown of *RS31* in the *i-rsT* lines caused an additional and pronounced reduction of the AS ratio, revealing that *RS31* functions as a major negative regulator of *RS31a* expression. The quantitative change was slightly more pronounced for *i-rsT-II* than *i-rsT-I*, being in agreement with a stronger reduction of *RS31* transcript levels due to a more efficient action of the corresponding *amiR* in *i-rsT-II* (Fig. 5A). We also determined the relative levels of the productive *RS31a* variant to rule out that the AS ratio change resulted primarily from a decrease of the unproductive variants. We observed an up to ∼7-fold increase of the protein-coding *RS31a.1* variant when comparing the mock- and estradiol-treated samples for the *i-rsT* lines (Fig. 5C, right), implying that the level of the RS31a protein can be strongly upregulated under these conditions.

To further examine the splicing regulatory function of the RS proteins, we analyzed the AS patterns of *RRC1* and *PPD2*, two genes that have previously been shown to undergo AS changes in etiolated seedlings upon exposure to light or sucrose (Hartmann et al., 2016; Saile et al., 2023). Comparable to our finding for the *RS31a* splicing patterns, we observed already in the mock-treated samples of *i-rsT* and *i-rsQ* consistently lower ratios of the unproductive variant *RRC1.2* relative to the protein-coding variant *RRC1.1* in comparison to the WT, and a further decrease was seen upon estradiol-induced *RS31* knockdown (Fig. 5D). Quantification of the relative levels of *RRC1.1* confirmed the upregulation of the protein-coding variant in the mock-treated *i-rsT* and *i-rsQ* (Supplementary Fig. S10A), and showed a tendency of a slight additional induction upon *RS31* knockdown. Accordingly, the *RS* genes act as negative regulators of *RRC1,* which has been identified as positive regulator of photomorphogenesis (Shikata et al., 2012). Analysing the AS pattern of *RRC1* in the single *rs* knockout mutants and *rs31 rs31a_C_* (Supplementary Fig. S10B, C) revealed for all of them a slight ratio change, suggesting that all four *RS* genes contribute to negative regulation of *RRC1*. We also investigated AS of *RRC1* upon treating etiolated seedlings for 6 h with sucrose. In line with previous findings, the AS ratio *RRC1.2/.1* was strongly reduced by this treatment (Fig. 5D). Again, inducible knockdown of *RS31* caused a significant reduction in the ratio *RRC1.2/.1*, further demonstrating the role of *RS31* in regulating *RRC1* and highlighting that the treatment with sucrose alone had not resulted in a saturated response. Analysing from the same samples the AS patterns of *PPD2* demonstrated the function of *RS31* in the regulation of another light-responsive AS event (Supplementary Fig. S10D-F). In this case, the corresponding AS change was less pronounced and only significant upon estradiol treatment for the *i-rsT-II* and *i-rsQ-II* lines containing the more efficient *amiR-II*. It can be concluded from these data that AS of *PPD2* is regulated not only by the RS proteins, but also additional splicing regulators. In summary, our experiments showed that the RS proteins are involved in regulating light-responsive AS events. Knocking down *RS31* released its negative regulation from *RS31a* and *RRC1*, both of which can support photomorphogenesis and may contribute to the reduced hypocotyl elongation observed under these conditions.

## Discussion

### *RS* genes are integrated in a cross-regulatory network of RBPs

Our experiments revealed extensive negative feedback regulation within the subfamily of *RS* genes. Accordingly, knocking out *RS40* and *RS41* caused an upregulation of *RS31* transcript levels and a relative decrease of unproductive transcripts for *RS31a*. The inducible knockdown of *RS31* in the *rs40 r41_C_* background resulted in an additional and pronounced decrease in the ratio of unproductive transcripts to the protein-coding variant of *RS31a*, with the levels of the latter variant being ∼7-fold increased in comparison to the WT. Analysis of previous RNA-seq data revealed that total transcript levels of *RS31*, *RS40*, and *RS41* were in a similar range, while *RS31a* abundance was ∼10-fold lower. Accordingly, *RS31a* is normally strongly repressed by the other *RS* genes, and this repression is released in the higher order *rs* mutant. Negative cross-regulation of *RS31a* by the other *RS* genes is the consequence of an AS shift towards unproductive variants containing PTCs that can trigger turnover via nonsense-mediated decay (NMD). This coupling of AS and NMD (AS-NMD) is a common principle among various mechanisms of negative feedback control, which is involved in fine-tuning the expression of a large number of RBPs (Müller-McNicoll et al., 2019). The prevalence of AS-NMD and its importance for the regulation of the entire family of *SR* genes has been described first for human and mouse (Lareau et al., 2007b; Lareau et al., 2007a) and was then also systematically examined in Arabidopsis (Palusa and Reddy, 2010), including the detection of NMD-sensitive AS variants from all four *RS* genes. Nuclear retention of specific AS variants represents another means to prevent their translation and has also been reported for *SR* genes from Arabidopsis, e.g., for an intron-retaining *RS2Z33* transcript variant (Göhring et al., 2014), an *RS31* splicing variant (Petrillo et al., 2014), and an *SR30* splicing isoform that is formed upon usage of a light-repressed alternative 3’ splice site (Hartmann et al., 2018). Further experiments are required to fully understand the auto- and cross-regulation that the *RS* genes are subjected to, both within their subfamily and in the context of the other *SR* genes. Earlier studies indicated that *SR30* could act as a negative regulator of *RS31*, based on an increase in unproductive *RS31* splicing variants in *SR30* overexpression lines (Lopato et al., 1999). Moreover, Kalyna et al. (2006) reported that AS of *RS31* is not auto-regulated. Based on iCLIP experiments from a recent preprint, RS31 can bind to the transcripts of all four *RS* genes (Köster et al., 2024). In line with our finding of negative cross-regulation of *RS31a* by *RS31*, they observed a decrease in unproductive splicing for *RS31a* and in addition for *RS40* in an *rs31* T-DNA insertion line.

Regulating closely related RBP genes via negative auto- and cross-regulatory feedback loops allows balancing their expression and achieving a robust output. Well-studied examples from Arabidopsis include the *GLYCINE-RICH RNA-BINDING PROTEIN* genes *GRP7* and *GRP8* (Staiger et al., 2003; Schöning et al., 2007; Schöning et al., 2008) and *POLYPYRIMIDINE TRACT BINDING PROTEIN* genes *PTB1* and *PTB2* (Stauffer et al., 2010; Rühl et al., 2012). Analyzing the AS patterns of single *ptb1* and *ptb2* mutants as well as the corresponding double mutant provided evidence for redundant and specific functions of these two splicing regulators (Rühl et al., 2012). The observation that the knockdown of *RS31* can induce photomorphogenesis compared to the negative effect seen upon loss of the other *RS* genes suggests that this cross-regulatory network serves in balancing the activity of these genes with at least partially antagonistic and not only redundant functions. This is further supported by our observation that the *RS* genes trigger AS-NMD and thereby repress *RRC1*, a positive regulator of phyB-dependent light responses (Shikata et al., 2012). Remarkably, the AS patterns of three *RS* genes was shown to be altered in an allelic series of T-DNA-carrying *rrc1* mutants, with a relative increase of the non-productive variants for *RS31*, *RS31a*, and *RS40* in seedlings grown under continuous light (Shikata et al., 2012).

Accordingly, *RRC1* acts as a positive regulator of its repressors. Moreover, *SFPS* was shown to be required for unproductive splicing of *RRC1* (Xin et al., 2019), representing another negative regulatory loop connecting these two photomorphogenesis-promoting factors. In a later study, SWAP1 was identified as an additional component of a ternary complex with RRC1 and SFPS functioning in light-responsive AS and promoting photomorphogenesis (Kathare et al., 2022). Given its categorization as RBP, it will be interesting to test whether SWAP1 can also influence AS of *RRC1*, *SFPS*, and the *RS* genes, which might further expand the intricate network of cross-regulation between RBPs involved in light-dependent AS and seedling development (Fig. 6).

**Fig. 6.**
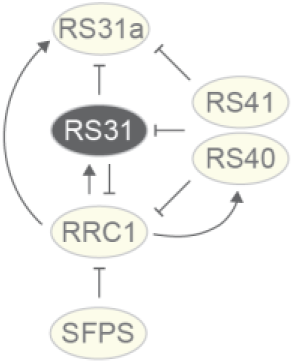
Model of cross-regulation between splicing regulators involved in light responses. Regulators of light-responsive splicing that promote (yellow ellipses) or suppress (RS31, black ellipse) photomorphogenesis form a network of positive (arrows) and negative (lines with bar) interactions.

### Physiological functions of *RS* genes

Based on the extent of hypocotyl elongation in response to red light, *RS40*, *RS41*, and *RS31a* act as activators of photomorphogenesis. An opposite function of *RS31* can be concluded from the observation that its knockdown caused a pronounced reduction in hypocotyl length in the background of mutants lacking two or three of the other *RS* genes. Thus, the network of *RS* genes can function in stabilizing the physiological response to light. Cotyledon opening represents another key parameter of light-dependent seedling development, and all *RS* overexpression lines displayed increased opening angles compared to the WT in darkness, with the effect being most pronounced in the *RS41* overexpression line. This line showed also reduced hypocotyl elongation, particularly in response to red light, whereas the other *RS* overexpression lines had WT-like hypocotyl lengths. Whether the distinct phenotype upon overexpressing *RS41* compared to the other *RS* genes is linked to specific features of *RS41* or the extent of overexpression remains to be examined. In agreement with our findings, Martín et al. (2025) recently reported in a preprint a positive role of RS40 and in particular RS41 in cotyledon opening, which they further linked to ABA-dependent splicing regulation.

Interestingly, we observed that overexpression of *RS31* also slightly increased cotyledon opening in darkness, indicating its functioning in partial suppression of this skotomorphogenic feature. Given that the hypocotyls were shorter upon *RS31* downregulation, it is supposed to act in this process as a suppressor of photomorphogenesis. These partially opposing effects regarding skoto-/photomorphogenesis might be explained in different ways. First, the elevated *RS* levels in the overexpression lines could trigger common AS shifts or other changes in gene expression that result in increased cotyledon opening in darkness. Consequently, the specificity of individual *RS* genes might be lost upon its overaccumulation, unlike under normal conditions when their expression is balanced within an RBP network. Second, reduced hypocotyl elongation upon *RS31* knockdown was observed in the background of a double and triple knockout of the other *RS* genes, and their absence might also contribute to this phenotype. Third, the *RS* genes might have different activities and functions in the processes of hypocotyl elongation and cotyledon opening.

Scoring development over the whole plant life cycle revealed additional critical functions of the *RS* genes. Simultaneous loss of *RS40* and *RS41* resulted in the generation of shortened siliques, while the *rs* quadruple mutant almost completely lost fertility due to a pollen defect. Interestingly, a triple mutant lacking besides *RS40* and *RS41* also *RS31a* had the same silique length as the *rs40 rs41_C_* mutant, suggesting that *RS31a* does not play an additional role and cannot partially compensate the loss of the two other *RS* genes in this developmental feature. This was different for hypocotyl elongation, as a more pronounced increase was seen in the triple compared to the double mutant. The additional loss of *RS31* in case of the quadruple mutant resulted in sterility, while seed generation was only slightly diminished in the double and triple mutants, highlighting the functional redundancy of the *RS* genes in this developmental process. The previous report of an *rs* quadruple mutant without any visible phenotype might be explained by residual activity of the respective mutant alleles carrying T-DNA insertions (Yan et al., 2017). Finally, the overall delayed development we observed upon overexpression of *RS31*, *RS40*, and *RS41*, as well as the dwarfism and leaf curling for the *RS41* over-accumulating line pointed at additional developmental aspects that can be regulated by the *RS* genes. It seems likely that these processes also involve networks of *RS* and other RBP genes, as described in this work in the course of light-dependent seedling development.

### Light-regulated AS in plant development

This work describes the critical role of the *RS* genes in light-dependent seedling development and demonstrates their involvement in the regulation of the light-responsive AS events within *RS31*, *RRC1*, and *PPD2*. Previous reports linking RS function with AS include a change in a heat stress-regulated exitron event upon *RS31* overexpression (Cecchini et al., 2022) and increased intron retention for several genes in an *rs40 rs41* mutant (Chen et al., 2013). Transcriptome-wide analyses of the AS patterns in the various *rs* misexpression lines established in the course of this work will allow to define the *RS* genes’ global impact on the large number of AS events that change in a light-dependent manner in etiolated seedlings (Shikata et al., 2014; Hartmann et al., 2016) but also at other developmental stages such as seed germination (Tognacca et al., 2019). Furthermore, it will be of interest to obtain a better understanding how these splicing changes elicit the physiological adaptations during de-etiolation. Our current knowledge of the functional impact of light-induced AS events is limited to a few examples of light signaling genes, including downregulation of the negative factor *SPA1-RELATED 3* (Shikata et al., 2014) and upregulation of the positive regulator *RRC1* (Hartmann et al., 2016). Recent studies suggested that AS-mediated gene regulation is more common among key signaling components of light responses. Dong et al. (2020) described a phyB-dependent intron retention event within the 5’ UTR of the *PHYTOCHROME-INTERACTING FACTOR3 (PIF3)*, where the retained sequence can suppress translation of the main open reading frame and thereby may control PIF3 levels in a diurnal manner. AS of a rice homolog of *Elongated hypocotyl 5* (*HY5*), a positive regulator of photomorphogenesis, was reported to generate two protein-coding isoforms that might have specific functions in skoto- and photomorphogenesis (Bhatnagar et al., 2023).

AS-mediated gene control can enable rapid responses and nuclear detainment of intron-retaining variants was recently demonstrated to be a common mechanism preventing translation of the corresponding mRNAs in the light response in Arabidopsis (Zhou et al., 2024). The same study screened for phenotype suppressors of the *cop1-6* mutant, which is impaired in the function of the central photomorphogenesis repressor COP1 (CONSTITUTIVE PHOTOMORPHOGENIC 1). This resulted in the identification of the spliceosomal components *PRP8* (*PRE-MRNA PROCESSING 8*), *SLU7*, *CACTIN* and genes from the family of DEAD-box helicases. A large number of light-induced AS events were shown to depend on COP1, implying it can alter spliceosome activity in a light-dependent manner. In line with this, Li et al. (2022) reported the function of COP1 and the RNA helicase U2AF65-ASSOCIATED PROTEIN (UAP56) as regulators of AS and repressors of photomorphogenesis. Moreover, a defect in the methylosome complex, which is required for proper spliceosome functioning, affected not only splicing but also photomorphogenesis, leading to red and blue light hypersensitive in the hypocotyl length response (Mateos et al., 2023). Among the first splicing factors linked to light-dependent AS was SFPS (Xin et al., 2017), which co-localizes with U2 components of the spliceosome. Following studies could demonstrate that SFPS interacts with RRC1 (Xin et al., 2019) and SWAP1 (Kathare et al., 2022), resulting in a ternary complex that is supposed to control light-dependent AS. Interestingly, while SFPS, RRC1, and SWAP1 all can interact with phyB and function in red light-dependent signaling, a role of the multifunctional and RNA-related CCR4-NOT complex in phyA-mediated, far-red light-responsive AS was revealed (Schwenk et al., 2021). Besides these studies in Arabidopsis, regulators of red light-dependent intron retention were discovered from *P. patens*, including hnRNP-H1 (Shih et al., 2019) and hnRNP-F1 (Lin et al., 2020). Red light-triggered intron retention and phototropism in *P. patens* were also linked to histone methylation (Wang et al., 2021), adding another level of regulation. In summary, our identification of the RS proteins as regulators of light-regulated AS further expands the network of factors underlying this widespread phenomenon. Integrating the available data and further experiments, such as investigating the AS patterns in higher order mutants, will be required to clarify whether all of these factors contribute to a common AS response to light or if different light responses with potentially distinct functions might exist.

### Phosphorylation and localization of RS proteins

One particularly interesting and largely unsolved question is how light and metabolic signals can change the activity of the involved splicing factors and regulators to elicit the corresponding AS shifts. The work by Zhou et al. (2024) suggested that the composition and activity of the spliceosome might be directly regulated in a COP1-dependent manner.

Furthermore, the AS regulators SFPS (Xin et al., 2017), RRC1 (Xin et al., 2019), and SWAP1 (Kathare et al., 2022) were all shown to interact with phyB and co-localize with it in light-induced photobodies. Both the interactions and the subcellular localization could change their activity in AS regulation, and it remains to be addressed whether the proteins localized within the photobodies are available to directly influence the splicing process.

Interestingly, our localization studies in *N. benthamiana* did not provide evidence for RS protein accumulation in photobodies. While further experiments are needed to validate this finding in stably transformed plants, the distinct localization patterns might point at the existence of different regulatory mechanism. For example, recruitment of the splicing regulatory proteins to photobodies could result in the sequestration, modification, or degradation of the corresponding proteins. In line with previous studies showing RS localization in nuclear speckles (e.g. Docquier et al., 2004; Lorković et al., 2004; Tillemans et al., 2005), we observed RS41-GFP accumulation in such subnuclear domains for some cells. Interestingly, a more homogenous distribution within the nucleoplasm could also be seen in other cells, and further experiments are needed to examine the underlying mechanism for this variation and its potential functional implications. Earlier studies had shown that phospho-signaling can affect the subnuclear distribution of RS proteins.

Specifically, chemical inhibition of phosphatase activity with okadaic acid caused in case of RS31 an increase in speckle size and a decrease of the diffuse signal within the nucleoplasm (Tillemans et al., 2006; Tillemans et al., 2005; Docquier et al., 2004). Based on our observation of increased RS41 phosphorylation upon light and sugar exposure, it is tempting to speculate that the phosphorylation status might alter RS activity by altering its distribution within the nucleus. Extensive phosphorylation of RBPs including RS proteins has been demonstrated previously (van Fuente Bentem et al., 2006) and is also reflected by database entries such as in the plant PTM viewer (Willems et al., 2019). However, our current understanding of how phospho-signaling contributes to AS regulation is scarce, and we know little about the involved kinases and the functional impact of the phosphorylation of their target proteins (Rodriguez Gallo and Uhrig, 2023). Scarpin et al. (2020) observed dephosphorylation of RS40 and RS41 upon TOR inactivation, which might relate to the slightly increased levels of a transgene-derived RS41 protein observed in this study.

Moreover, a mutant deficient in members from the SR protein-specific kinase II family showed reduced phosphorylation and altered sub-cellular localization of RS and other SR/SR-like proteins (Wang et al., 2023), making these kinases interesting candidates for further studies. Unravelling the dynamics of RS protein phosphorylation and its impact on their function, including the subnuclear distribution in response to light and sugar signals, will be key aspects of future research to define the role of these enigmatic splicing regulators in seedling photomorphogenesis. The existence of four RS proteins with both redundant and specific sequence characteristics as well as physiological functions can provide a paradigm to gain a better understanding how RBP functions have evolved and diversified in plants.

## Experimental procedures

### Plant material and growth conditions

*Arabidopsis thaliana* (Arabidopsis) mutants generated in this study include the following CRISPR mutants: *rs31_c_*, *rs31a_c_*, *rs40_c_*, *rs41_c_*, *rs31 rs31a_c_*, *rs40 rs41_c_*, *rsT* (*rs31a rs40 rs41_c_*), *rsQ* (*rs31 rs31a rs40 rs41_c_*), *i-rsT* (*i-amiR rs31 rs40* rs41*_c_*) and *i-rsQ* (*i-amiR-rs31 rs31a rs40 rs41_c_*). All mutants are in the Columbia-0 (Col-0) background.

Arabidopsis seeds were surface sterilized using 3.75% sodium hypochlorite and 0.01% Triton X-100 and plated on ½ Murashige and Skoog (MS) medium (M0222.0050, Duchefa) pH 5.7 to 5.8, containing 0.8% (w/v) plant agar (Duchefa). Depending on the experiment, MS media contained 5 µM β-estradiol (E2758-1G; Sigma-Aldrich) or an equivalent concentration of dimethyl sulfoxide (DMSO; mock). After 2 d of stratification, germination was induced by illuminating the seeds with white light (∼100 μmol m^-2^ s ^-1^) for 2 h. Depending on the experiment, plates were either transferred to light or darkness.

For phenotyping experiments plants were grown on soil under long day conditions (16 h light, 8 h dark, 22 °C) with a regular light intensity (∼100 μmol m^-2^ s ^-1^).

### Generation of *rs* CRISPR mutants

For the assembly of two sgRNA expression cassettes, sgRNA inserts were amplified from pCBC-DT1T2 (Xing et al., 2014) using Phusion® High-Fidelity DNA Polymerase and two inner and two outer primers that were partially overlapping and contained the sgRNA target sites. pHEE401E vector (Wang et al., 2015) and sgRNA expression cassettes (T1T2-PCR) were purified and used for *Bsa*I cut ligation. The reaction was incubated in a thermocycler at 37 °C for 5 h, followed by 50 °C for 5 min and 80 °C for 10 min. Final pHEE401E_T1T2_RS-sgRNA constructs were transformed into *Agrobacterium tumefaciens* strain C58C1 and used for transforming Arabidopsis Col-0 wild-type plants via floral dipping (Clough and Bent, 1998). Seeds from T0 plants were surface sterilized using 80% ethanol and 0.05% Triton X-100 and plated on ½ MS containing 20 μg/ml hygromycin. Resistant seedlings (T1) were transferred to soil and genomic DNA was extracted to analyze mutations in corresponding *RS* genes. Therefore, fragments surrounding the target sites were amplified, purified, and sequenced. Transgene free plants, still containing the corresponding mutations, were identified in the subsequent generations. All oligonucleotides to generate CRISRP constructs and genotyping are listed in Supplementary Tab. S1. *rsT_c_* and *rsQ_c_* mutants were generated by transforming transgene free *rs40 rs41_c_* double mutant plants with *Agrobacterium tumefaciens* containing the *RS31 RS31a* CRISPR construct.

### amiRNA plasmid constructions and generation of transgenic plants

Two independent amiRNA sequences targeting *RS31* were identified and corresponding primers designed using the WDM3 Web microRNA Designer (WDM3, http://wmd3.weigelworld.org; Schwab et al., 2006; Ossowski et al., 2008). Site-directed mutagenesis on the endogenous miR319a precursor was performed by overlap PCR. To this end, the PCR was performed in three single PCR reactions using Q5® High-Fidelity DNA Polymerase (NEB), pRS300 as template and the following primer combinations: For *i-amiR- RS31-I* TW80/JS260, JS259/JS258 and JS257/TW81, for *i-amiR-RS31-II* TW80/JS264, JS263/JS262 and JS261/TW81. Purified PCR products were mixed with the outer primer pair TW80/TW81 and an overlap PCR for each construct was performed. Corresponding *amiR* precursor sequences were then transferred into the cloning vector pJET1.2 and used to assemble intermediate vectors via GreenGate reaction, resulting in pGGNJS01 and pGGNJS02. GreenGate reaction was performed as described in Lampropoulos et al. (2013). Finally, amiR intermediate vectors were combined with the estradiol-inducible intermediate vector that expresses XVE (pGGMTW01) (Saile et al., 2023) and the destination vector pGGZ003. Corresponding GreenGate reactions resulted in the final constructs pGGZJS01 and pGGZJS02. All modules used for the generation of the intermediate and destination vectors are listed in Supplementary Tab. S1.

Final constructs were transformed into *Agrobacterium tumefaciens* strain ASE, and dipped into Arabidopsis *rs40 rs41_c_* and *rsT_c_* (*rs31a rs40 rs41_c_*) CRISPR mutants, respectively, to generate an inducible triple and quadruple mutant. Transgenic plants were selected using 40 μg/ml Hygromycin.

### Generation of *RS* overexpression constructs and lines

Coding sequences (cds) of *RS31*, *RS31a*, *RS40* and *RS41* genes were cloned into the vector pBinAR-HA_3_ containing the constitutive CaMV promoter (Höfgen and Willmitzer, 1992). All cds were amplified from plasmids, with ES232/233 (*RS31*), AWTU374/375 (*RS31a*), AWTU376/377 (*RS40*) and ES248/249 (*RS41*). Sequences of oligonucleotides are listed in Supplementary Tab. S1. Generated inserts were cloned into pBinAR-HA_3_ via *Kpn*I/*BamH*I. Cloning of the construct for the vector control line *35S::EGFP* was previously described (Wachter et al., 2007). The constructs were transformed into Arabidopsis Col-0 via floral dipping (Clough and Bent, 1998).

### Generation of RS41-HA_3_ *tor* and *snrk1* transgenic plants, protein isolation, SDS-PAGE and Western blot

The 35S::RS41-HA_3_ construct was transformed into *i-amiR-SnRK1-I* and *i-amiR-SnRK1-II* mutant plants (Saile et al., 2023) via floral dipping and transgenic seedlings were selected by 25 μg/ml Kanamycin. The *i-amiR-TOR* construct (pAP44) (Saile et al., 2023) was dipped into transgenic plants overexpressing RS41-HA_3_ and transgenic seedlings were selected by 3 mM D-Alanine.

To analyze RS41 protein level in *tor* and *snkr1* mutant background, etiolated seedlings were grown in liquid ½ MS media. On the third day, etiolated seedlings were either treated with 5 μM β-estradiol or DMSO, serving as corresponding mock control. Subsequently, seedlings were grown for further 3 d under dark conditions, followed by a treatment with either 2% sucrose or white light (∼100 μmol m^-2^ s^-1^) for the indicated time points. 0.1 g of 6-d-old seedlings were flash frozen, ground in liquid nitrogen and resuspended in 0.1 ml extraction buffer (50 mM Tris-HCl pH 7.5, 200 mM NaCl, 4 M Urea (w/v), 0.1% (v/v) Triton X-100, 5 mM DTT, 1x protease inhibitor cocktail). Lysates were clarified by centrifugation at 10,000g and 4 °C for 10 min and proteins denatured at 70 °C for 10 min in SDS sample buffer. Proteins were separated by 10% SDS-PAGE and transferred to a nitrocellulose membrane using wet transfer (110 V, 400 mA, 1 h, 4 °C). Membranes were blocked in 5% skim milk for 1 h, followed by the first antibody incubation overnight at 4 °C. Primary antibodies α-HA (1:2000) and α-tubulin (1:10,000) were combined. Secondary antibody incubation was performed for 1 h at room temperature and chemiluminescence was imaged using the Fusion Fx system (Vilber). Relative band intensities were quantified using the quantification tool of the Evolution-Capt Edge program, followed by calculating the volume ratio of HA to Tubulin.

Commercial antibodies were used as followed: rat α-HA (11867423001, Roche), rabbit α-tubulin (AS10680, Agrisera). Anti-rat (31470, Thermo Fisher Scientific) and anti-rabbit (A6154, Sigma-Aldrich) secondary antibodies were conjugated with horseradish peroxidase.

### Cotyledon opening assay

Seedlings were grown horizontally on ½ MS plates in darkness for 6 d. On the sixth day, etiolated seedlings were carefully transferred under green light to ½ MS plates supplemented with 1.5% plant agar (pH 5.7 – 5.8). Plates were scanned and cotyledon opening angle was measured using ImageJ. Cotyledons with angles that were greater than 6 degrees were counted as opened.

### Hypocotyl assay

To measure hypocotyl length, seeds were stratified for 2 d at 4 °C. Germination was induced by exposing the seeds to regular white light (∼100 µmol m^−2^ s^−1^) for 2 h before placing them in darkness or under the corresponding light quality. Seedlings were grown horizontally on ½ MS plates in darkness or continuous monochromatic red (peak: ∼655 nm), blue (peak: 447/456 nm), or far-red (peak: ∼730 nm) light with varying intensities for 4 d. Depending on the experiment, ½ MS plates were additionally supplemented with 5 µM β-estradiol or DMSO as control. To reach decreasing light intensities in the same growth chamber, plates were sealed with black tape and lids were covered with an increasing number of white paper layers. On the fourth day, etiolated seedlings were transferred to new ½ MS plates and scanned. Hypocotyl length was measured using the ImageJ software.

### Light and sucrose treatment

Seedlings were grown in liquid ½ MS media (pH 5.7 – 5.8) under dark conditions. On the third day, 5 μM β-estradiol or DMSO was added to the liquid media under green light conditions, followed by a growth for further 3 d in darkness. On the sixth day, MS media was exchanged by liquid ½ MS media supplemented with either 1.06% mannitol or 2% sucrose. The new media also contained either 5 μM β-estradiol or DMSO. Subsequently, seedlings were either kept in darkness or exposed to white light (∼100 μmol m^-2^ s^-1^) for 6 h.

### RNA extraction, RT-qPCR, and PCR product analyses

Total RNA was isolated from Arabidopsis seedlings using the Universal RNA Purification Kit (Roboklon, EURx), with an on-column DNaseI treatment. Reverse transcriptions were performed using either Superscript II Reverse Transcriptase (Invitrogen) or AMV Reverse Transcriptase Native (Roboklon, EURx), according to the manufacturer’s instructions. RT-qPCR was conducted using MESA GREEN qPCR Mastermix on a CFX384 real-time PCR cycler (Bio-Rad) and data were analyzed as previously described (Stauffer et al., 2010). Co-amplification PCRs were performed using homemade Taq polymerase, visualized by agarose gel electrophoresis and quantified on a Bioanalyzer using the DNA1000 kit (Agilent Technologies). Corresponding primer sequences are listed in Supplementary Tab. S1.

### *In vivo* localization studies in *N. benthamiana*

GoldenGate cloning was used to generate the pUbi::RS41-GFP construct. First, the *RS41* cds was amplified from plasmid DNA using Q5® High-Fidelity DNA Polymerase (NEB) with primers JS190/JS191. The resulting PCR product was then A-tailed and ligated into the pGEM-T Easy vector (Promega). To remove internal *Bsa*I and *Bpi*I restriction sites, site-directed mutagenesis was performed using the QuickChange Lightning Multi Site-Directed Mutagenesis Kit (Agilent) with primers JS299/JS300, following the manufacturer’s protocol. Mutations were confirmed by sequencing, and the modified construct, along with other modules, was used to assemble the destination vector via a *Bsa*I GoldenGate reaction. The GoldenGate reaction was incubated under the following conditions: 37 °C for 5 min and 16 °C for 5 min, cycled 25 times, followed by 5 min at 50 °C and 5 min at 80 °C. This reaction resulted in the final construct, BB10_JS03. A list of all primers and modules can be found in Supplementary Tab. S1. For co-localization studies with PHYB-tagRFP, the pCHF230-PHYB vector was cloned. pCHF230 is a T-DNA vector containing a *p35S:BamHI-XbaI- tagRFP:terRbcS* cassette and *BlpR* for selection using BASTA, and was obtained as follows: The *tagRFP* cds was PCR amplified from pFRETvr-2in1-NN (Hecker et al., 2015) using primers AH1/AH2. The resulting PCR fragment was digested with *Bam*HI and *Spe*I and ligated into pCHF5 (Hiltbrunner et al., 2005) opened with *Bam*HI and *Xba*I, resulting in the vector pCHF230. The pCHF230-PHYB vector, containing a *p35S:PHYB-tagRFP:terRbcS* cassette, was constructed as follows: The *PHYB* cds was excised from pCHF40-PHYB (Sheerin et al., 2015) using *Xba*I and ligated in sense orientation into pCHF230, which had been linearized with *Xba*I. Successful cloning was confirmed by sequencing the insert and flanking regions.

Agrobacteria-mediated leaf infiltration of *N. benthamiana* was performed according to (Wachter et al., 2007). *Agrobacterium tumefaciens* containing constructs *pUbi::RS41-GFP* and *p35S::PHYB-tagRFP* were grown overnight at 28 °C. The next day, Agrobacteria cultures were centrifuged, cells resuspended in water, and OD_600 nm_ adjusted to 0.8. Infiltration mixtures were prepared by mixing equal amounts of both constructs and a P19 construct for silencing suppression. Subsequently, Agrobacteria mixtures were infiltrated in leaves of 3- to 4-week-old *N. benthamiana* plants. Three days after infiltration, leaf discs were imaged using a Leica TCS-SP5 Laser Scanning Confocal Microscope using photomultiplier tube (PMT) detectors and a 63x glycerol objective. GFP was detected using an argon laser with excitation wavelength of 488 nm and collection range of 498 - 535 nm. RFP and autofluorescence were detected using a DPSS laser with excitation wavelength of 561 nm and collection range of 571 - 695nm.

### Statistical analyses

If not otherwise stated, statistical analyses have been performed using GraphPad Prism 9 or R 4.3.3. The results are provided in Supplemental Data Set S2.

### Phosphoproteomics

#### Plant material and growth conditions

Arabidopsis Col-0 seedlings were grown in liquid ½ MS media in darkness for 6 d. On the sixth day, media was exchanged with either ½ MS media or ½ MS media supplemented with 2% sucrose. Subsequently, plates were kept in darkness (i. dark, ii. dark and sucrose treatment) or shifted to regular white light (iii. light treatment, ∼100 μmol m^-2^ s^-1^). After 30 min, seedlings were harvested and flash frozen.

#### Protein extraction and precipitation

2 to 3 g plant material was ground in liquid nitrogen and suspended in extraction buffer (8 M urea, 40 mM NaCl, 50 mM Tris pH 7.5, 2 mM MgCl_2_, PhosSTOP (Roche), phosphatase inhibitor mix I (Serva), Complete Protease Inhibitor Mixture (Roche)) in a 1:4 ratio. Cell debris were collected by centrifugation (15 min, 16,000g, 4 °C) and supernatant was filtered through Miracloth. Proteins were precipitated using methanol/chloroform. To the supernatant, 4 volumes of cold methanol, 1 volume of cold chloroform and 3 volumes of cold milliQ water was added, with mixing by vortexing in between. Lysates were centrifuged at 16,000g for 5 min, 4 °C. Top layer was discarded and 4 volumes of cold methanol added to the lower and interphase to further precipitate the protein layer. Samples were centrifuged again (16,000g for 5 min, 4 °C) and supernatant discarded afterwards. Precipitated protein was washed with cold methanol and centrifuged at 16,000g for 5 min, 4 °C. Washing steps were repeated four times and precipitated protein pellets air dried.

#### In solution digest

Chloroform-Methanol precipitated samples were dissolved in 6 M urea/2 M thiourea (Sigma) buffered in 10 mM HEPES (pH 8.0) rotating at 1000 rpm for 30 min. Dithiothreitol (Thermo Fisher) was added to a final concentration of 1 mM and incubated at room temperature (RT) for 45 min rotating at 300 rpm. Proteins were alkylated with a final concentration of 5.5 mM chloroacetamide (Sigma) at RT for 1 hour in the dark with gentle shaking. To digest the proteins, trypsin (Sigma) in a 1:150 ratio was added for an overnight incubation at RT. The digestion was stopped by pipetting trifluoroacetic acid (TFA, final 0.5%, Fisher Chemicals) to the samples and by incubating at 4 °C for at least 30 minutes. Peptides were centrifuged at 4 °C and 3500g for 10 minutes and purified on a Sep-Pak Vac 6cc C18 500 mg Cartridges (Waters). Columns were activated before with 5 ml 100% acetonitrile (ACN) and washed three times with 5 ml 0.1% TFA. After loading the clarified peptide mixture, the column was washed again three times with 10 ml 0.1 % TFA. Peptides were eluted with 4.5 ml 50% acetonitrile (ACN, Sigma) and the concentration measured with a NanoDrop spectrometer (A_280_ value).

#### Phosphopeptide enrichment

The eluate (diluted with TFA to a final concentration of 6%) was incubated with Titansphere TiO_2_ (GL Science) in a 1.5-fold ratio TiO_2_ beads to peptides for 1 h on a rotation wheel. The peptide-bound beads were washed two times with 50% ACN, 6% TFA and again two times with 50% ACN, 0.1% TFA with a 1000g centrifugation step in between the washes. Beads were transferred to C8 47 mm extraction disk (CDS Analytical), assembled in a 200 µl pipette tip and centrifuged at 600g. Enriched phospho-peptides were eluted in two steps (5% ammonia and 10% ammonia, 25% ACN) at 400g in a 2 ml tube. The eluate was vacuum-concentrated at 45 °C for 15 min and acidified with TFA.

#### C18 stage tipping

For desalting and peptide concentration C18 tips were prepared by cutting out two C18 47 mm extraction disks (CDS Analytical) and placing them into a 200 µl pipette tip. The C18 tips were equilibrated with 25 μl methanol, followed by 25 μl stage tip buffer B (80% ACN, 0.1% formic acid (Sigma) and 2 x 25 μl stage tip buffer A’ (3% ACN, 1% TFA) by centrifuging at 500g. The samples were loaded on the tip and washed with 50 μl buffer A (0.1 %FA). The peptides were eluted with 50 μl elution buffer (50% ACN, 0.1% formic acid) in a 24-well Thermo-Fast PCR-plate (Thermo Fisher) and vacuum-concentrated at 45 °C to a volume of 4 to 5 µl. For the further mass spectrometry analysis, the sample was diluted with buffer A to 6 to 7 µl.

#### Mass spectrometer data acquisition

Peptide fractions were analyzed on quadrupole Orbitrap mass spectrometers (Exploris 480) equipped with a UHPLC system (EASY-nLC 1200, Thermo Scientific) (Bekker-Jensen et al., 2020). Samples were loaded onto C18 reversed-phase columns (length 55 cm, inner diameter 75 μm; 1.9 μm C18 bead size) and eluted with a linear gradient from 2.4 – 32% ACN containing 0.1% FA in 2 h. The mass spectrometer was operated in data-dependent mode, automatically switching between MS and MS^2^ acquisitions. MS spectra (m/z 325–1300) and fragment spectra were acquired in the Orbitrap mass analyzer. The 20 most intense ions were sequentially isolated and fragmented by HCD (Olsen et al., 2007). Peptides with unassigned charge states, as well as with charge states less than +2 were excluded from fragmentation.

#### MS analysis

Raw data files were analyzed using MaxQuant (version 2.3.0.0) (Cox and Mann, 2008). Parent ion and MS2 spectra were searched against the database Araport11 (released in 2016) using Andromeda search engine (Cox et al., 2011). Spectra were searched with a mass tolerance of 6 p.p.m in MS mode, 20 p.p.m. in HCD MS2 mode, strict trypsin specificity, and allowing up to two mis-cleavages. Cysteine carbamidomethylation was searched as fixed, while N-terminal acetylation and methionine oxidation were searched as variable modifications. Match between runs was activated. The dataset was adjusted to a 1% false discovery rate on a peptide spectrum match level based on a target-decoy approach using reversed protein sequences.

Statistical analysis and data filtering was performed using Perseus software v2.0.9.0 (Tyanova et al., 2016). Proteins “reverse hits”, or “potential contaminants” were removed. Only proteins identified with at least a localization probability > 0.75 were used for the analysis. Peptides were further filtered for at least two valid values in at least one of the treatment groups. To identify significantly enriched proteins, a p-value-controlled t-test was performed. The p-value cut-off was set to 5% and s0 to 0.1.

## Supporting information

Supplemental Data Set 1

Supplemental Data Set 2

## Funding

This project was funded by the Deutsche Forschungsgemeinschaft (DFG, German Research Foundation) - SFB 1551 – Project No. 464588647. This research project was further supported by the DFG with grants to AW (Project Nos. 232631280: SFB1101/C03 and 270473865). During the writing phase of the manuscript, JS was affiliated with and funded by the Laboratory of Plant Physiology, Wageningen University & Research, 6708 PB Wageningen, The Netherlands.

## Author contributions

Conceptualization: AW and JS; Investigation: JS, HW, PL, MD, LS, KL, and SG; Resource contribution: AH; Writing: AW, JS and HW; Supervision: AW and PB; Funding acquisition: AW and PB.

## Supplementary data

**Supplementary Figure S1**. Phospho-proteome analysis upon light and dark treatment of etiolated seedlings.

**Supplementary Figure S2**. RS41 protein level is increased upon TOR and SnRK1 knockdown.

**Supplementary Figure S3**. Light regulation of RS gene expression. **Supplementary Figure S4**. Knocking out all *RS* genes results in male sterility. **Supplementary Figure S5**. Screening of *i-rsT* and *i-rsQ* mutants.

**Supplementary Figure S6**. Hypocotyl elongation of higher order *rs* mutants under red, far-red, and blue light.

**Supplementary Figure S7**. RS41-GFP does not accumulate in phyB-containing photobodies.

**Supplementary Figure S8**. Impact of *RS* overexpression on plant development.

**Supplementary Figure S9**. Hypocotyl lengths in *RS* overexpression lines.

**Supplementary Figure S10**. Altered splicing patterns in *rs* mutants.

**Supplementary Table S1**. List of oligonucleotides and constructs.

**Supplementary Data Set S1**. Phospho-peptide data of light- and sucrose treated seedlings.

**Supplementary Data Set S2**. Statistical analyses.

## Acknowledgements

We are grateful to Xu Na Wu and Waltraud X. Schulze for sharing unpublished data of phospho-proteome experiments with us. Furthermore, we thank Claudia König and Lara Cronhardt-Lück-Gießen for technical assistance.

## Data statement

The MS-based proteomics data have been deposited with the ProteomeXchange Consortium via the PRIDE partner repository with the dataset identifier PXD062307. Any additional information required to re-analyze the data reported in this paper is available from the lead contact upon request.

## Notes

### Competing Interest Statement

The authors have declared no competing interest.

