## Supplemental Figures for "A network of RS splicing regulatory proteins controls light-dependent splicing and seedling development"

**Saile *et al.*, Supplemental data**

**A**

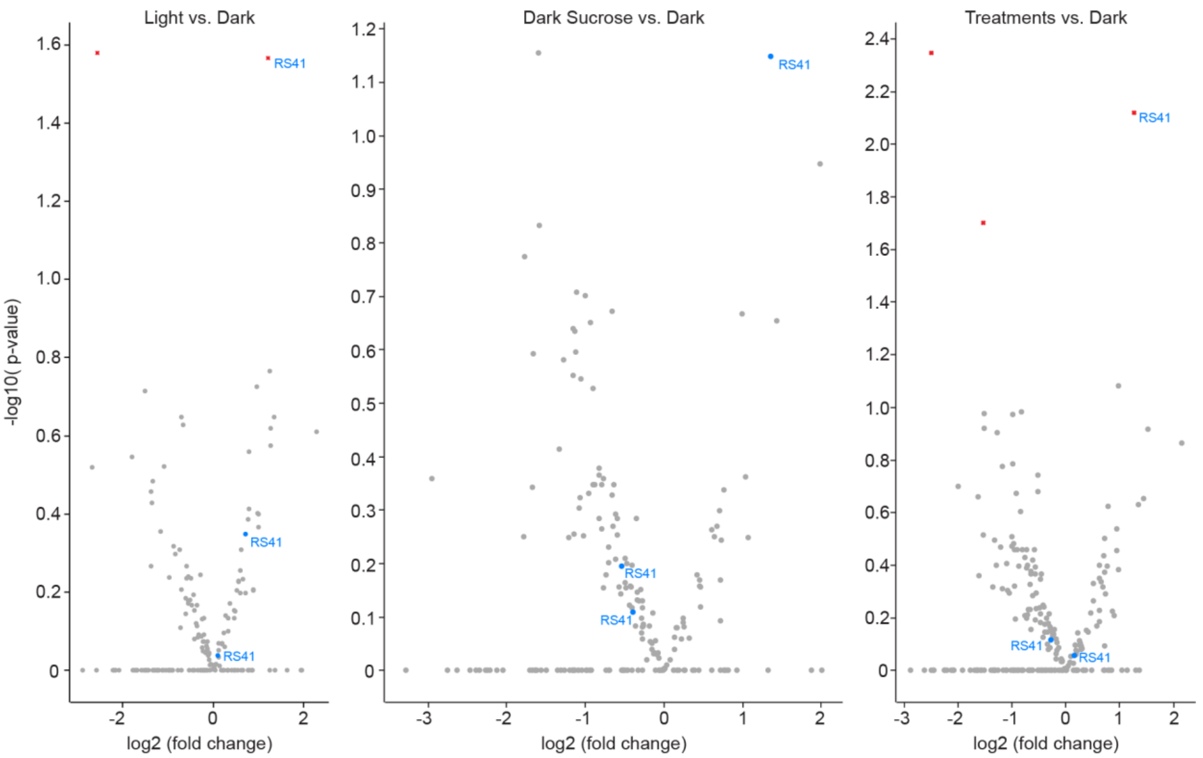


**B**

**RS31 (AT3G61860.1, P92964-1), 264 aa**

MRPVFVGNFEYETRQSDLERLFDKYGRVDRVDMKSGYAFVYFEDERDAEDAIRKLDNFPFGYEKRRLSVEWAKGE
RGRPRGDAKAPSNLKPTKTLFVINFDPIRTKEHDIEKHFEPYGKVTNVRIRRNFSFVQFETQEDATKALEATQRS
KILDRVVSVEYALKDDDERDDRNGGRSPRRSLSPVYRRRPSPDYGRRPSPGQGRRPSPDYGRARSPEYDRYKGPA
AYERRRSPDYGRRSSDYGRQRSPGYDRYRSRSPVPRGRP

**RS31a (AT2G46610.1, Q9ZPX8-1), 250 aa**

MRHVYVGNFDYDTRHSDLERLFSKFGRVKRVDMKSGYAFVYFEDERDAEDAIRRTDNTTFGYGRRKLSVEWAKDF
QGERGKPRDGKAVSNQRPTKTLFVINFDPIRTRERDMERHFEPYGKVLNVRMRRNFAFVQFATQEDATKALDSTH
NSKLLDKVVSVEYALREAGEREDRYAGSRRRRSPSPVYRRRPSPDYTRRRSPEYDRYKGPAPYERRKSPDYGRRS
SDYGRARARSPGYDRSRSPIQRARG

**RS40 (AT4G25500.1, P92965-1), 350 aa**

MKPVFCGNFEYDAREGDLERLFRKYGKVERVDMKAGFAFVYMEDERDAEDAIRALDRFEFGRKGRRLRVEWTKSE
RGGDKRSGGGSRRSSSSMRPSKTLFVINFDADNTRTRDLEKHFEPYGKIVNVRIRRNFAFIQYEAQEDATRALDA
SNNSKLMDKVISVEYAVKDDDARGNGHSPERRRDRSPERRRRSPSPYKRERGSPDYGRGASPVAAYRKERTSPDY
GRRRSPSPYKKSRRGSPEYGRDRRGNDSPRRRERVASPTKYSRSPNNKRERMSPNHSPFKKESPRNGVGEVESPI
ERRERSRSSPENGQVESPGSIGRRDSDGGYDGAESPMQKSRSPRSPPADE

**RS41 (AT5G52040.1, P92966-1), 356 aa**

MKPVFCGNFEYDARESDLERLFRKYGKVERVDMKAGFAFVYMEDERDAEDAIRALDRFEYGRTGRRLRVEWTKND
RGGAGRSGGSRRSSSGLRPSKTLFVINFDAQNTRTRDLERHFEPYGKIVNVRIRRNFAFIQYEAQEDATRALDAT
NSSKLMDKVISVEYAVKDDDSRGNGYSPERRRDRSPDRRRRSPSPYRRERGSPDYGRGASPVAHKRERTSPDYGR
GRRSPSPYKRARLSPDYKRDDRRRERVASPENGAVRNRSPRKGRGESRSPPPYEKRRESRSPPPYEKRRESRSPP
PYEKRRERSRSRSKSSPENGQVESPGQIMEVEAGRGYDGADSPIRESPSRSPPAEE

**SFig. 1. Phospho-proteome analysis upon light and dark treatment of etiolated seedlings. A)** Scatter plots depicting phosphorylation differences in 6 d-old etiolated seedlings that were exposed for 30 min to light (left) or to 2% sucrose (middle) in comparison to seedlings kept in darkness and control medium, and scatter plot showing analysis of combined data from the two treatments relative to the control (right). Significant changes are depicted by red symbols; RS41 phospho-peptides are labelled accordingly. **B)** Sequences of RS proteins with the RNA recognition domains being underlined. Serine (S), threonine (T), and tyrosine (Y) residues highlighted with blue boxes were reported to be phosphorylated according to the database from the plant PTM viewer (Willems et al., 2019). Serine residues in blue letters were found to be phosphorylated upon light and sucrose treatment of etiolated seedlings in at least one of two phospho-proteome experiments.

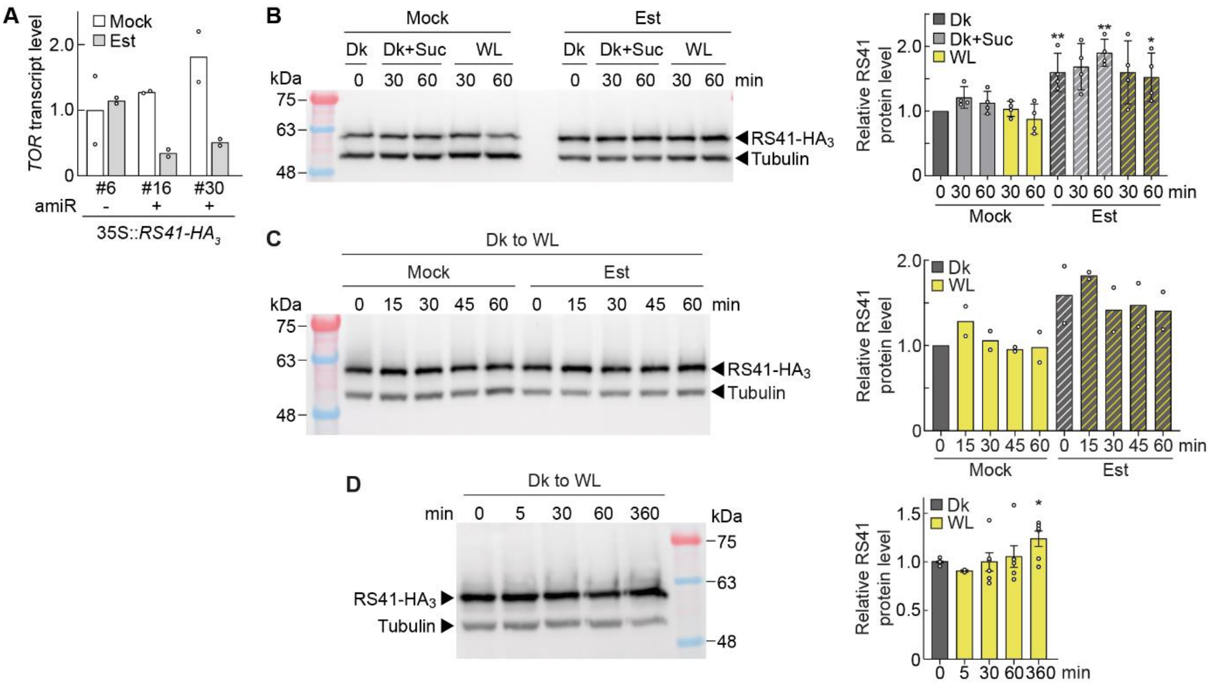


**SFig. 2. RS41 protein level is increased upon TOR and SnRK1 knockdown. A)** Relative transcript level of *TOR* in 6-d-old etiolated *RS41* OE seedlings in a background without (-) or with (+) an *i-amiR*-construct, treated with either mock (white bars) or estradiol (Est, grey bars) for 3 d before sampling. Data are mean values (n = 2) and individual data points are shown as dots. Numbers below the bars refer to individual lines. **B)** (Left) Representative immunoblot detection of RS41-$HA_3$ and Tubulin in 6-d-old *i-amiR-TOR* #16 seedlings. Dark-grown seedlings were treated with either mock or estradiol for 3 d, followed by a sucrose treatment in darkness (Dk+Suc) or white light exposure (WL) for 30 or 60 min. (Right) Quantification of RS41 protein level in *i-amiR-TOR* #16 and #30 seedlings. Bar graph depicts mean values (n = 4) of two replicates each from the two independent *i-amiR-TOR* mutant lines, based on the ratio of the RS41-$HA_3$/Tubulin signal and normalized to the mock control at 0 min. Statistical significance was determined by two-tailed Student's t-test against corresponding mock control (P values: *P < 0.05, **P < 0.01). Details on statistical testing for this and following figures provided in Supplemental Data Set S2. **C)** (Left) Representative immunoblot of RS41-$HA_3$ and Tubulin in 6-d-old etiolated *i-amiR-SnRK1* seedlings containing a *35S::RS41-HA*$_3$ construct. *i-amiR-SnRK1* seedlings were previously established (Saile et al. 2023) and transformed with the *35S::RS41-HA*$_3$ construct. 3-d-old etiolated seedlings were treated with either Mock or estradiol for 3 d. At day 6, seedlings were illuminated with white light (WL) for the indicated time points. (Right) Quantification of RS41/Tubulin protein ratio in *i-amiR-SnRK1* seedlings. Data are mean values (n = 2), normalized to the mock control at 0 min. Individual data points are shown as dots. **D)** (Left) Representative immunoblot detection of RS41-$HA_3$ and Tubulin as loading control in 6-d-old seedlings carrying the *35S::RS41-HA*$_3$ construct in a WT background. Other details as described for B) and C). Quantification is based on 2 – 6 replicates represented by circles. Statistical significance was determined by two-tailed Student's t-test against dark grown (0 min) sample (P value: *P < 0.05).

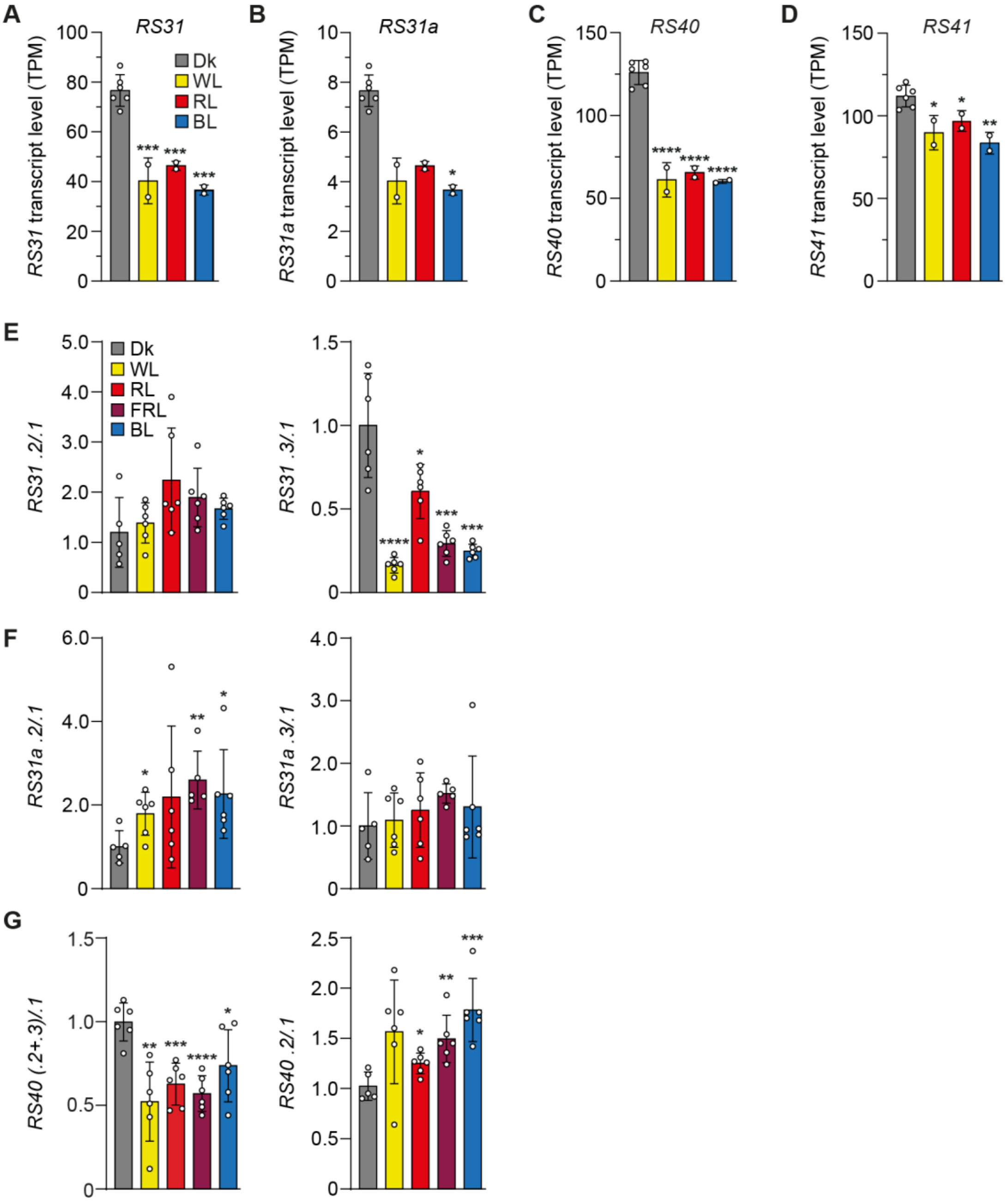


**SFig. 3. Light regulation of *RS* gene expression. A – D)** Gene expression levels of *RS31* (A), *RS31a* (B), *RS40* (C), and *RS41* (D) in 6-d-old etiolated WT seedlings that were illuminated with white light (WL; ~130 µmol m$^{-2}$ s$^{-1}$), red light (RL; ~14 µmol m$^{-2}$ s$^{-1}$), or blue light (BL; ~6 µmol m$^{-2}$ s$^{-1}$) for 6 h or kept in darkness (Dk), respectively. Expression levels are based on RNA-seq data from Hartmann et al. (2016). The dark samples are the control samples for each light quality and were combined in this figure, depicted as mean values ± SD (n = 6). Asterisks indicate significant differences compared to dark sample based on two-tailed t-test (P values: *P < 0.05, **P < 0.01, ***P < 0.001, ****P < 0.0001). **E – G)** Splice isoform ratios of *RS31*, *RS31a,* and *RS40* in 6-d-old etiolated WT seedlings that were either kept in darkness (Dk) or exposed to white light (WL, ~10 µmol m$^{-2}$ s$^{-1}$), red light (RL, ~8 - 11 µmol m$^{-2}$ s$^{-1}$), far-red light (FRL; ~6 – 11 µmol m$^{-2}$ s$^{-1}$), or blue light (BL~8 -11 µmol m$^{-2}$ s$^{-1}$) for 6 h. Data represents mean ± SD, normalized to the mean dark control, with individual data points shown as dots (n = 5 to 6). A two-tailed student's t-test was performed to analyse significant differences compared to the dark control (P values: *P < 0.05, **P < 0.01, ***P < 0.001, ****P < 0.0001).

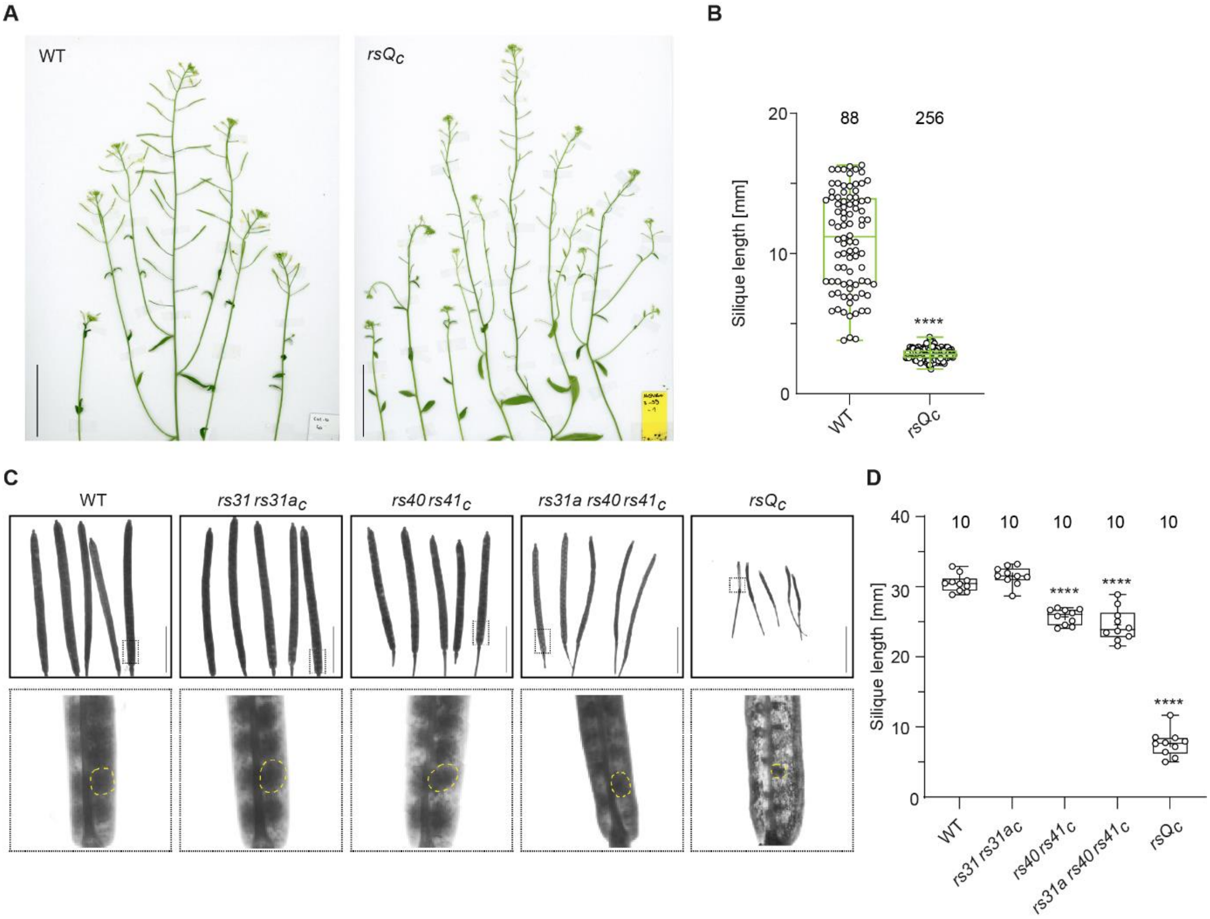


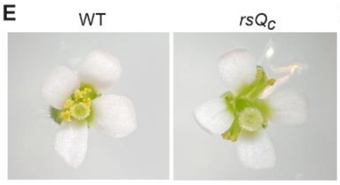


F

| Female | x | Male | n[1] | n[2] | generated siliques [%] |
|---|---|---|---|---|---|
| *rsQc* | | WT | 73 | 67 | 92 |
| WT | | *rsQc* | 26 | 1 | 4 |

[1]total pollinated flowers, [2]total generated siliques

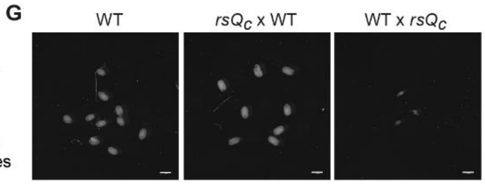


**SFig. 4. Knocking out all *RS* genes results in male sterility. A)** Representative scans of 43 d-old WT and *rsQc* mutant inflorescences. Plants were grown under long day conditions. Scale bar: 5 cm. **B)** Quantification of silique length at the age of 43 d. Median is represented by the central line; box limits show the 25th and 75th percentiles, and whiskers go down to the smallest value and up to the largest. n is indicated at the top. Asterisks indicate significant differences compared to WT based on two-tailed t-test (P value: ****P < 0.0001.) **C)** (Upper panel) Representative pictures of siliques from WT and different *rs* mutant plants at the age of 50 d. Scale bar: 1 cm. (Lower panel) Magnification of the boxed regions displayed in the upper panel, with one seed per genotype marked by a dashed sphere. **D)** Quantification of silique length from plants described in C. Asterisks indicate significant differences compared to WT based on one-way ANOVA with post-hoc Tukey test (P value: ****P < 0.0001). **E)** Photographs of representative WT and *rsQc* flower, showing mis-formed anthers and pollen defects in *rsQc*. **F)** Generation of siliques from a crossing experiment of *rsQc* mutant plants that were pollinated with WT pollen or vice versa. **G)** Representative pictures of seeds derived from pollination experiment. WT seeds derived from self-pollination served as control. Scale bar: 1 mm.

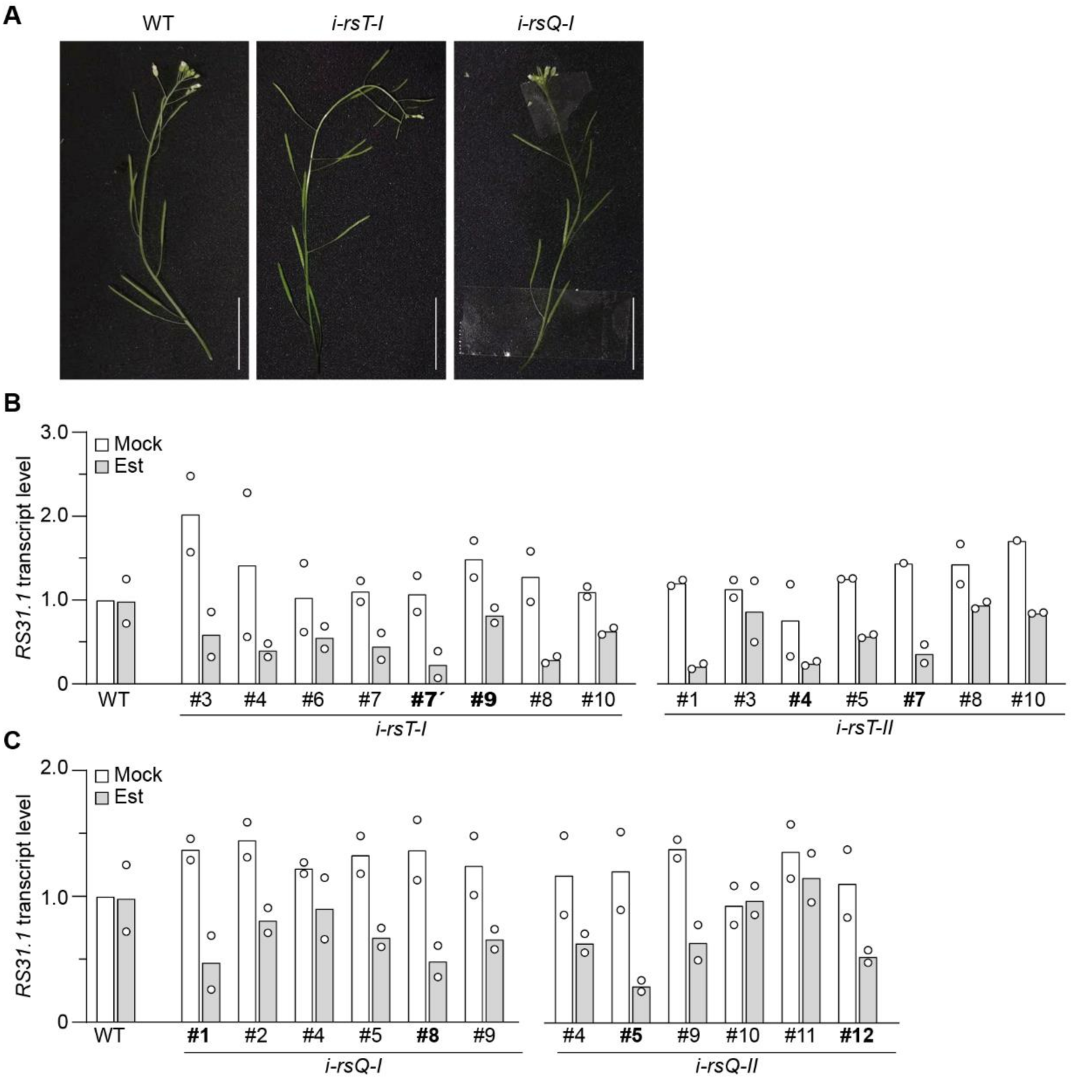


**SFig. 5. Screening of *i-rsT* and *i-rsQ* mutants. A)** Representative scans of 7-week-old WT, *i-rsT-I,* and *i-rsQ-I* inflorescence stems with siliques. Plants were grown in the absence of estradiol. Scale bar: 1 cm. **B, C)** Relative transcript level of *RS31.1* in comparison to WT Mock in *i-rsT* (**B**) and *i-rsQ* (**C**) mutants. Two independent amiR constructs (**I** and **II**) were generated, both for targeting *RS31*. Seedlings were grown for 6 d in darkness and treated with either mock (white bars) or estradiol (Est, grey bars) at the third day. Data are mean values (n = 2; individual data points shown as dots). For each line, a pool of 10 sublines was used. Lines with names in bold were used for further experiments.

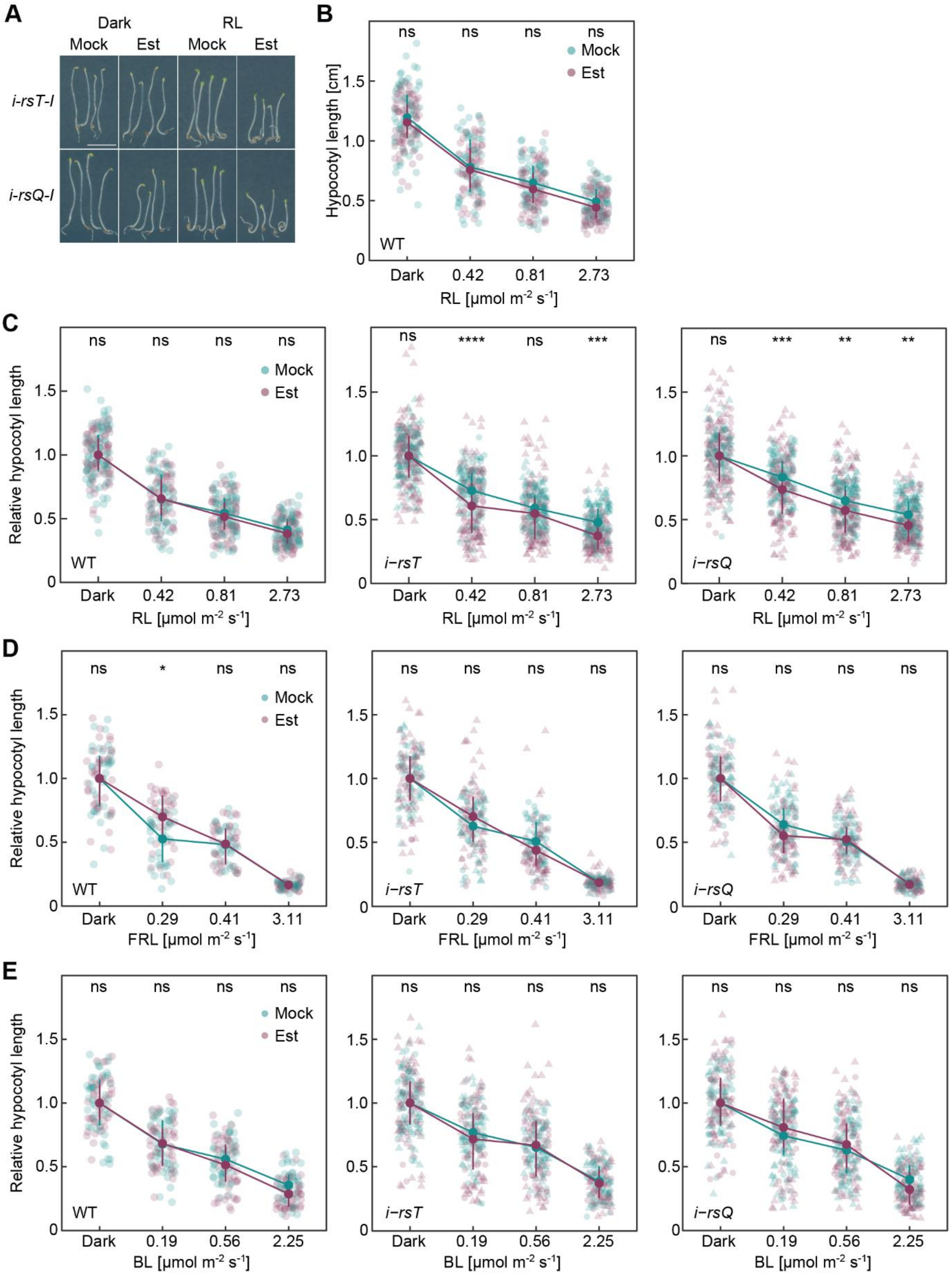


**SFig. 6. Hypocotyl elongation of higher order *rs* mutants under red, far-red, and blue light. A)** Representative pictures of *i-rsT-I* (*i-amiR-RS31-I* in *rs40 rs41*$_C$) and *i-rsQ-I* (*i-amiR-RS31-I* in *rs31a rs40 rs41*$_C$) seedlings grown for 4 d in darkness or red light (RL, 0.42 µmol m$^{-2}$ s$^{-1}$) under mock or Estradiol (Est) treatment. White scale bar corresponds to 0.5 cm. **B)** Absolute hypocotyl length of WT grown

under indicated light qualities and intensities or in darkness on plates containing either mock or Est. Interquartile range and mean are depicted as vertical line and circle, respectively, and asterisks indicate significant differences of the comparison between Est and mock samples based on Kruskal-Wallis test with Dunn's *post-hoc* test (α = 0.05) and ns > 0.05 (n = 56 to 82; individual data points shown as dots). **C – E)** Relative hypocotyl length of WT, *i-rsT*, and *i-rsQ* grown under indicated RL (C), far-red light (FRL, D), and blue light (BL, E) intensities or in darkness on plates containing either mock or Est. Values were normalized to average length of dark-grown seedlings for each genotype. Interquartile range and mean are depicted as vertical line and circle, respectively. Asterisks indicate significant differences of the comparison between Est and mock samples based on Kruskal-Wallis test with Dunn's *post-hoc* test (α = 0.05) and ns > 0.05, *P ≤ 0.05, **P ≤ 0.01, ***P ≤ 0.001, ****P ≤ 0.0001 (n = 33 to 153; individual data points shown as dots). Circles and triangles indicate two lines with independent *amiR-RS31* constructs (I and II) which were used.

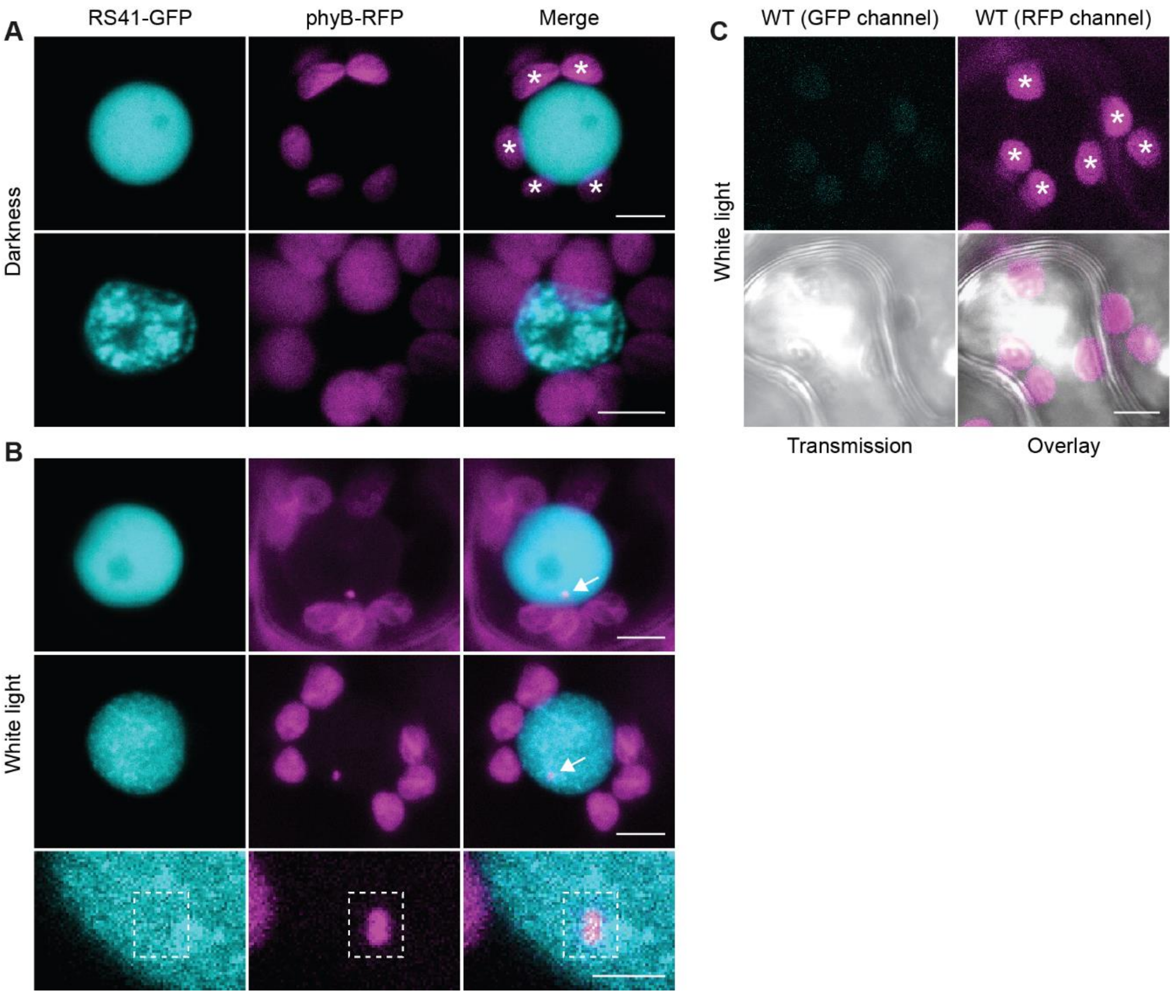


**SFig. 7. RS41-GFP does not accumulate in phyB-containing photobodies. A, B)** Representative images of RS41-GFP and phyB-RFP which were co-expressed in *N. benthamiana*. After infiltration, plants were either transferred to darkness (A) or kept in white light (B). Three days post infiltration, subcellular localisation was analysed using confocal microscopy. Arrows point to phyB-containing photobodies, which form in nuclei from light-incubated plants (B) but not in darkness (A). The last row in B shows a magnification of a region including the photobody. Scale bar in the magnification corresponds to 2 µm; all other scale bars depict 5 µm. Asterisks indicate chloroplasts showing autofluorescence in the RFP channel under dark and light conditions. **C)** Fluorescence signal of a non-transformed WT sample imaged with the same settings as for (A, B). Other display features as described before.

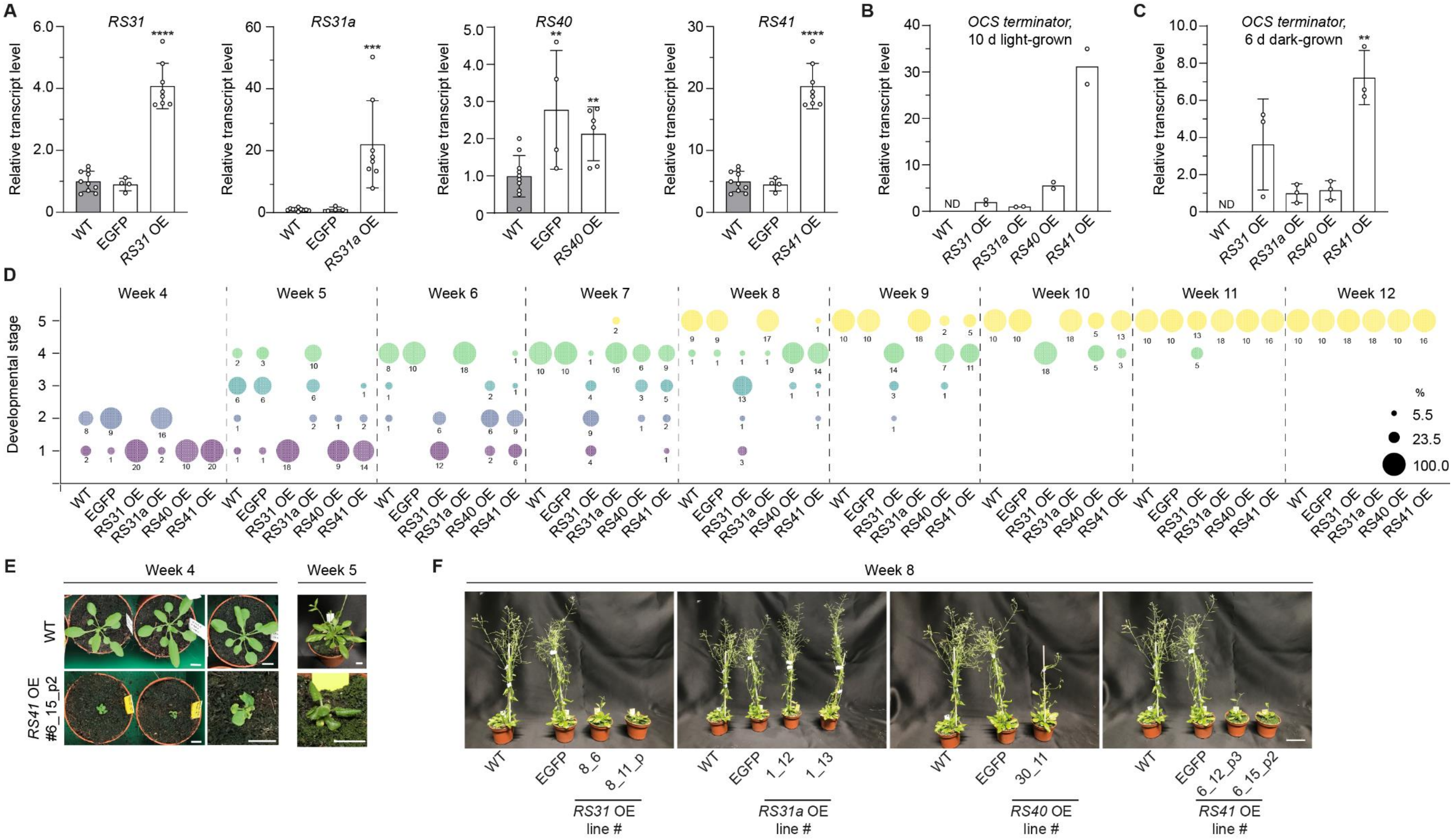

**SFig. 8. Impact of *RS* overexpression on plant development. A)** Relative transcript levels of the coding isoform of *RS31*, *RS31a*, *RS40,* and *RS41* in corresponding *RS* OE seedlings. WT (marked in grey) and a vector control line (*35S::EGFP*) were included as controls. All seedlings were grown in light for 10 d on ½ MS plates supplemented with kanamycin, except for WT that was grown on ½ MS plates w/o antibiotic. Data are mean values ± SD (n = 4 to 10,

individual data points shown as dots), that were normalized to the mean level in WT. Two corresponding *RS* OE sublines each were combined for the analysis as follows: For *RS31*, sublines from T2 (n = 4) and T3 (n = 4) were combined. For *RS31a*, both sublines were derived from T2 (each n = 4). For *RS40*, sublines from T2 (n = 4) and T3 (n = 2) were combined. For *RS41*, both sublines were derived from T3 (each n = 4). Statistical significance was determined by two-tailed Student's t-test against WT (P values: *P < 0.05, **P < 0.01, ***P < 0.001, ****P < 0.0001). **B, C)** Transgene expression based on RT-qPCR analysis with primers binding to the common *OCS* terminator region in 10-d-old seedlings (B), as described in (A) with n = 2 and 6-d-old etiolated *OE* seedlings (C) that were treated with 1.06% mannitol for 6 h (n = 3). Data was normalized to mean *OCS* transcript level in *RS31a* OE and values represent means (± SD in C). Individual data points are shown as dots. Statistical significance was determined by two-tailed Student's t-test against *RS31a* OE (P values: **P < 0.01; in C). **D)** Developmental stages of plants grown under long-day conditions and scored into the following stages once a week: 1, rosette; 2, bolting; 3, first opened flower; 4, no new buds; 5, (first) siliques visible. The size of each bubble is proportional to the percentage of plants at the corresponding stage of development and at the time shown. n = 8 to 20 plants per OE line and n = 10 per control line. Data of *RS31*, *RS31a,* and *RS41* are based on two sublines with 8 to 10 plants per subline. For *RS31*, two sublines from T2 and T3 were combined; for *RS31a,* both combined sublines were derived from T2 and for *RS41* both from T3. **E)** Close up picture of representative 28-d-old (week 4) and 35-d-old (week 5) WT and *RS41* OE plants. All scale bars: 1 cm. **F)** Representative photographs of 50-d-old *RS* OE and control lines from the phenotyping experiment. Scale bar: 5 cm.

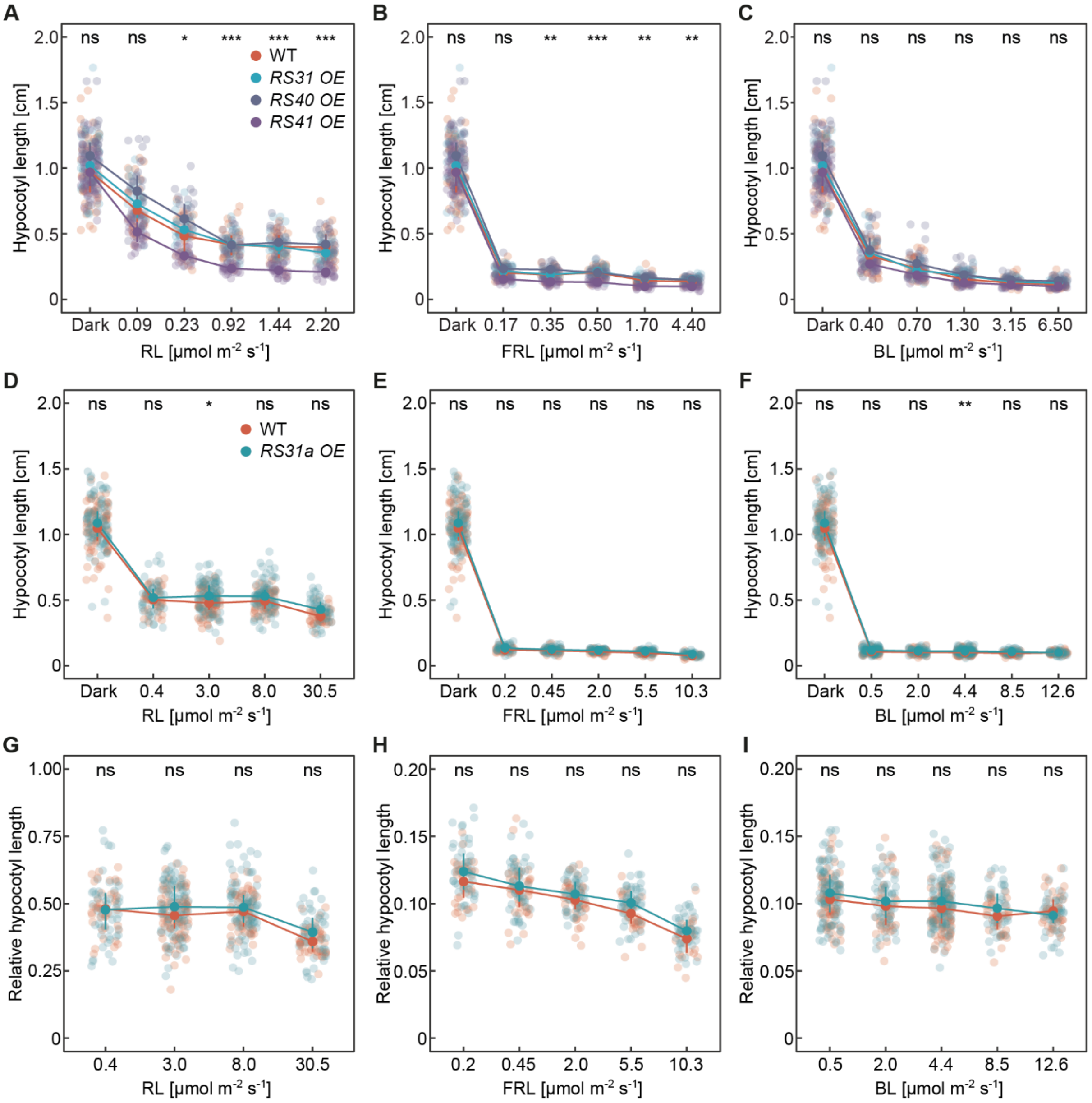


**SFig. 9. Hypocotyl lengths in *RS* overexpression lines. A - C)** Hypocotyl length of WT and indicated *RS* OE lines grown for 4 d on plates under indicated continuous red light (RL, A), far-red light (FRL, B), or blue light (BL, C) conditions. Interquartile range and mean are depicted as vertical line and circle, respectively. Asterisks indicate significant differences of the comparison between WT and *RS41 OE* based on Kruskal-Wallis test with Dunn's *post-hoc* test (α = 0.05) and ns > 0.05, *P ≤ 0.05, **P ≤ 0.01, ***P ≤ 0.001, ****P ≤ 0.0001 (n = 25 to 61; individual data points shown as dots). **D – I)** Absolute (D – F) and relative (G – I) hypocotyl lengths of WT and *RS31a OE* line grown under conditions and displayed as described before. Relative values resulted from normalization to average length of dark-grown seedlings for each genotype. Asterisks indicate significant differences of the comparison between the genotypes based on Kruskal-Wallis test with Dunn's *post-hoc* test (α = 0.05) and ns > 0.05, *P ≤ 0.05, **P ≤ 0.01 (n = 22 to 114; individual data points shown as dots).

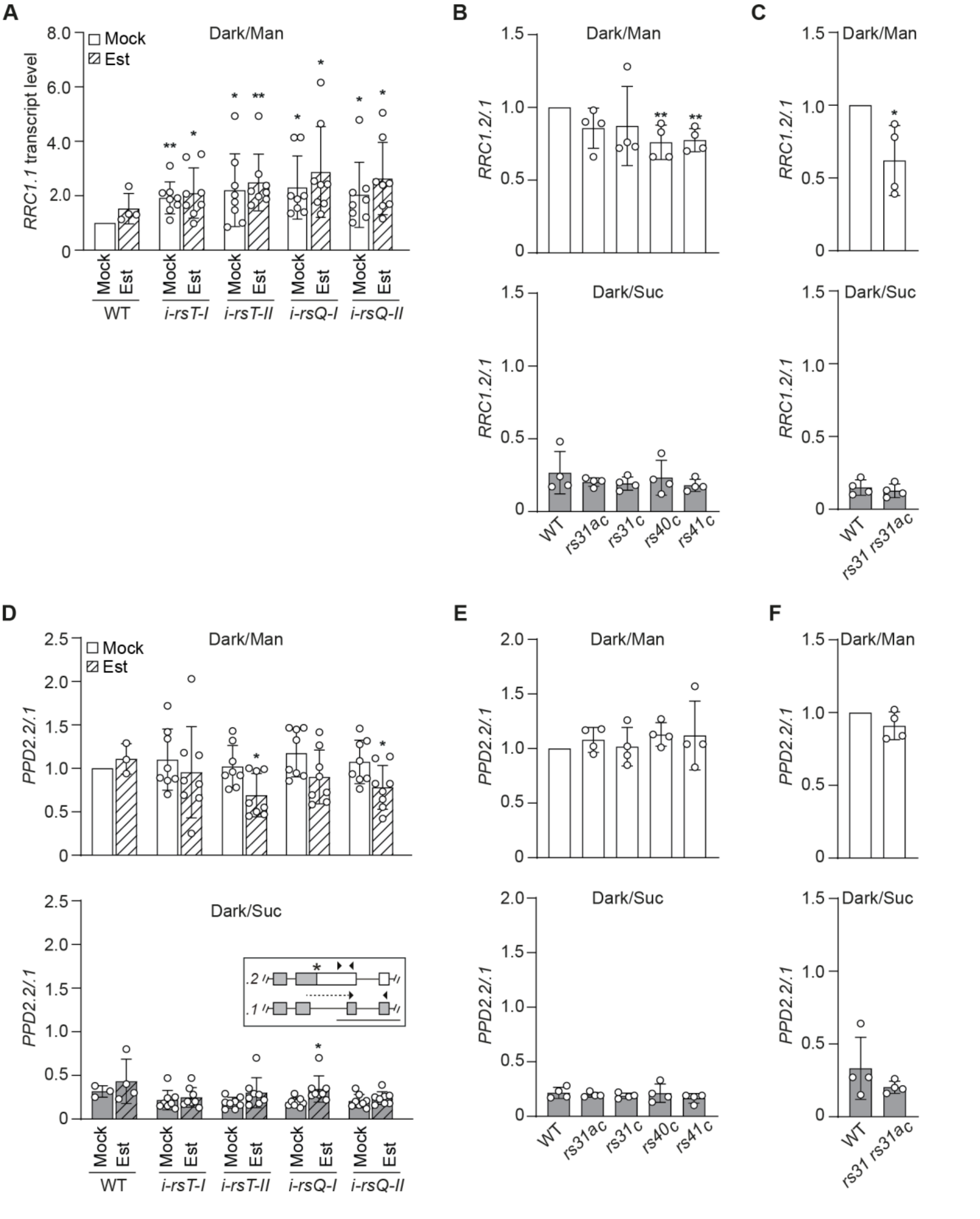


**SFig. 10. Altered splicing patterns in *rs* mutants. A)** Relative transcript level of *RRC1.1* in 6-d-old etiolated seedlings, treated with either mock (white bars) or estradiol (Est, hatched bars) for 3 d. Displayed are mean values (n = 4 – 8; individual data points are shown as dots) ± SD. Data was normalized to WT mock, dark + mannitol. Significant differences were analysed using a one-sample t-test, comparing values against the WT mock, which was set to 1 (P values: *P < 0.05, **P < 0.01). **B, C)** *RRC1* splicing pattern in 6-d-old seedlings, quantified via RT-qPCR. Seedlings were grown in liquid ½ MS media for 6 days under dark conditions. Statistical significance was determined by two-tailed Student's t-test against corresponding Mock control (P values: *P < 0.05, **P < 0.01). At day 6, media was exchanged with liquid ½ MS media, supplemented with either 1.06% mannitol (Man) or 2%

sucrose (Suc) and plates were kept in darkness for 6 h. Displayed are mean values (n = 4; individual data points shown as dots) ± SD, normalized to the WT dark + mannitol sample. **D - F)** *PPD2* splicing pattern in 6-d-old seedlings that was quantified via RT-qPCR. Growth conditions, treatments, and statistical analysis for (E, F) were as described in (B, C). For (D), etiolated seedlings were grown in liquid ½ MS media. After 3 d, estradiol (Est) and DMSO (mock) was added to the media, respectively. At 6 d, media was exchanged by liquid ½ MS media, supplemented with 1.06% mannitol or 2% sucrose, in the presence of estradiol and mock, respectively. Plates were further kept in darkness for 6 h. Displayed are mean values (n = 3 – 8, individual data points are shown as dots) ± SD, normalized to the WT mock, dark + mannitol sample. A two-tailed Student's t-test was performed in comparison to corresponding mock control (P values: *P < 0.05). *PPD2* splicing model is shown as inset, with lines representing introns, and grey and white boxes displaying UTRs and coding exons, respectively. Asterisk marks the position of a premature termination codon, and arrowheads depict binding sites of primers used for RT-qPCR. Scale bar: 500 nts.
